# Planting set-up and local environmental conditions affect outcomes in tropical agricultural restoration

**DOI:** 10.64898/2026.10.09.757530

**Authors:** Sacchi Shin-Clayton, Anak Agung Ketut Aryawan, Damayanti Buchori, Jean-Pierre Caliman, Purnama Hidayat, Eka Lupitasari, Mohammad Naim, Willow Oatley, Sarah H. Luke, Edgar C. Turner

## Abstract

1. Fine-scale environmental metrics, including spatial configuration and identity of surrounding plants, can strongly influence plant survival and ecosystem functioning. However, tropical restoration projects rarely test their role. Within the Riparian Ecosystem Restoration in Tropical Agriculture (RERTA) Project, located within oil palm plantations in Indonesia, we investigated how the spatial configuration and position of enrichment-planted trees influenced restoration outcomes within actively restored riparian margins.

2. We compared the enrichment plantings’ response to fine-scale spatial conditions, either planted with conspecific (same-species) or heterospecific (mixed-species) neighbours for six planted tree species *(Peronema canescens, Shorea leprosula, Alstonia scholaris, Artocarpus integer, Calophyllum inophyllum, and Albizia chinensis)*, in restoration areas with mature oil palms either left in place (shaded) or removed (unshaded). We also tested whether restoration outcomes varied with position within the margin (river, middle, or edge). We measured plantings response as seedling survival and height (monitored over eight years) for all six species of trees.

3. We also carried out more detailed observation on multiple ecosystem functions associated with focal trees of three of the species (*Peronema canescens*, *Alstonia scholaris*, and *Albizia chinensis),* measuring differences by planting configuration and distance to river edge. Functions measured included: herbivore predation, leaf herbivory, litterfall, soil chemical properties (organic carbon, cation exchange capacity, total nitrogen, and phosphorus), and soil physical properties (bulk density, aggregate stability, friable soil content, and A-horizon depth).

4. Survival and height did not differ between conspecific and heterospecific neighbours, but did with position within the riparian margin, but only when analysed by treatment and time. Planting configuration and position influenced ecosystem functions; heterospecific plantings had greater soil aggregate stability and higher ant densities, while increasing distance from the river was associated with lower arthropod abundance and herbivory but greater predation.

5. Our findings demonstrate that planting outcomes are strongly context-dependent, with local seedling configurations, position within the riparian margin, and restoration treatment interacting to influence restoration success and local ecological conditions. Our findings provide experimental results to inform planting strategies and restoration protocols in agricultural landscapes.

## Introduction

The location and spatial placement of plants within an environment have long been known to influence their survival, growth and resulting ecosystem processes (Comita et al., 2014; Fichtner et al., 2018). A key mechanism proposed to explain neighbourhood effects on vegetation is the Janzen–Connell hypothesis, which predicts that seeds and seedlings experience higher mortality near conspecific adults or at high conspecific densities because host-specific enemies, such as pathogens and herbivores, are more abundant there (Connell, 1971; Janzen, 1970) . This concept has been applied and tested in the five decades since it was proposed, with general support found for negative density- and distance-dependent seed and seedling survival in tropical forests (Comita et al., 2014; Webb & Peart, 1999). However, the scale and direction of these effects are impacted by species identity, the life stage of plants and environmental contexts (Comita et al., 2014; Zhu et al., 2015). The literature is also context-biased, with most tests conducted in naturally regenerating forest communities or through short-term seed or seedling experiments, with comparatively few publications focussed on later life stages or planted restoration communities (Comita et al., 2014; Ramage et al., 2017). There is less understanding of whether neighbourhood-dependent effects influence the performance of trees planted during restoration (Comita et al., 2014). This represents an important knowledge gap for tropical forest restoration, as this could heavily influence project success.

Enrichment planting, the planting of trees into degraded or regenerating landscapes, is increasingly used to accelerate forest recovery and re-establish species unable to recolonise naturally (Meli et al., 2026). However, outcomes vary considerably with environmental conditions, species identity and planting strategy (dos Santos et al., 2025; Jevon et al., 2022). A recent global synthesis identified only 74 enrichment- planting studies across 67 tropical and subtropical locations, most of which focused on tree survival and growth, with relatively few assessing wider ecosystem functions (Meli et al., 2026). Despite this limited evidence base, existing studies indicate that the local neighbourhood can influence planting success. Tropical tree seedlings planted beneath conspecific rather than heterospecific trees have been found to have lower survival and growth and greater foliar damage (Bagchi et al., 2010; Schweizer et al., 2013), while belowground associations with ectomycorrhizal communities may further facilitate responses to neighbour identity (Jevon et al., 2022; Schweizer et al., 2013). Together, these findings suggest that neighbouring tree identity and its associated biotic factors can influence restoration outcomes. However, whether these neighbourhood effects persist over longer timescales or change with time, extend beyond individual tree performance to wider ecosystem functions, or remain consistent across contrasting restoration environments remains poorly understood.

Understanding these processes is particularly important in tropical agriculture, where restoration increasingly occurs within degraded environments. Agricultural expansion has caused extensive tropical forest loss and habitat conversion (Gibbs et al., 2010; Pendrill et al., 2022). For example, oil palm expansion has contributed substantially to forest conversion in Indonesia and Malaysia (Xu et al., 2022). Within established oil palm systems, riparian areas along permanent waterways are increasingly recognised as important targets for protection and restoration (Barclay et al., 2017; Luke et al., 2020). However, restoration practitioners still lack fundamental knowledge about how planting configuration and the local environment influence restoration success, and how to restore these areas most effectively.

Few field experiments have tested how planting diversity and spatial configuration influence enrichment planting within oil palm plantations. An early project planted native trees along a river to establish a habitat corridor but did not manipulate planting configuration (Yamada et al., 2016). The EFForTS-BEE experiment advanced this approach by manipulating tree-island size and diversity, from monocultures to mixtures of two, three and six native species (Teuscher et al., 2016). After one year, island size, rather than planted diversity, influenced bird and invertebrate communities. Subsequent analyses extended this to local neighbourhood effects, examining tree–tree interactions, edge effects and competition with surrounding oil palms (Zemp, Ehbrecht, et al., 2019; Zemp, Gérard, et al., 2019). After five years, larger islands supported greater diversity and ecosystem functioning without reducing oil palm yield at the landscape scale (Zemp et al., 2023). However, these studies have largely manipulated diversity at the patch scale or examined neighbourhood effects arising within planted mixtures, rather than testing experimentally how the spatial arrangement of conspecific and heterospecific neighbours affects restoration outcomes under different environmental conditions.

Here, we address this knowledge gap using the Riparian Ecosystem Restoration in Tropical Agriculture (RERTA) experiment, located in oil palm plantations in Indonesia. In this experiment, we planted six native tree seedling species along river margins in different configurations, representing a landscape-scale field experiment that assesses the effects of conspecific or mixed-species neighbourhoods under contrasting restoration treatments (Luke et al., 2020). We test whether planting configuration influences long-term tree survival, height, and associated ecosystem functions, and whether these effects vary by restoration treatment (shaded or unshaded) and distance from the river. Specifically, we ask whether **(1)** seedlings planted in conspecific plantings experience reduced survival and height relative to those in mixed-species plantings, consistent with conspecific negative density dependence; **(2)** planting configuration influences associated ecosystem functions; and **(3)** these responses vary with distance from the river and between restoration treatments.

## Methods

### Study sites

The RERTA Project is based in industrial plantations in Riau province, central Sumatra, Indonesia (0°55′56″ N, 101°18′0″ E). It includes three alternative restoration treatments (each covering an area 400m long and 50m wide on either side of the river), replicated across two rivers, with treatments established during oil palm replanting (first-generation oil palm replanted with second-generation oil palm).

Treatments include: 1) remnant mature palms left in the margin and enrichment plantings with a mix of six native tree species (hereafter ‘Mature Palms and Plantings’), 2) remnant mature palms left in place and the margin left to regenerate naturally, 3) remnant mature palms removed and enrichment plantings with the same six native tree species (hereafter ‘Plantings Only’), 4) remnant mature palms removed and young oil palms replanted to the river’s edge (a negative control treatment).

For this study, we focus on fine-scale spatial comparisons within the actively restored margins only: Mature Palms and Planting Only. Within these two margins, an equal number of six native tree species (*Peronema canescens* (Sungkai); *Shorea leprosula* (Meranti tembaga); *Alstonia scholaris* (Pulai); *Artocarpus integer* (Cempedak); *Calophyllum inophyllum* (Bintangur); *Albizia chinensis* (Sengon)) were planted, which were selected from local nurseries and a range of functional traits (e.g. fruiting, leguminous, and pioneer species; Luke et al., 2020). Within both treatments, we carried out more detailed observations of ecosystem functioning for three of the species (*Peronema canescens*, *Albizia chinensis* and *Alstonia scholaris*), testing two planting configurations (hereafter, ‘planting type’): 1) seedlings were planted surrounded by the same species (hereafter, ‘conspecific’), and 2) seedlings were planted surrounded by different species from the focal species (hereafter, ‘mixed’). We also tested the effect of position within the riparian margin (river, middle, edge closest to the plantation).

### Enrichment planting field measurements

#### Seedling survival and height

We monitored the enrichment-planted seedlings for survival and height over eight years for all six tree species. Surveys were initially intended to occur approximately every six months; however, fieldwork was interrupted by the COVID-19 pandemic and subsequent changes in plantation management. Overall, a field team based at the Sinar Mas Agro Resources and Technology Research Institute (SMARTRI) Research Centre conducted nine surveys between 2017 and 2025, with margin area surveyed varying by year.

At each survey, seedlings were recorded as alive or dead and, where possible, their height was measured in centimetres. As trees became taller, height was measured using a clinometer and trigonometric approach, following FAO (2023a). Measurements were taken from a known horizontal distance from the tree, with the observer’s eye height accounted for when calculating total tree height.

#### Data processing and classification

##### General data processing

The long-term seedling monitoring dataset contained 54,223 observations across the nine surveys. We identified individual seedlings by combining river site, treatment, riverbank side, planting line, and within-line tree number to create a unique tree ID. This allowed us to link observations from the same seedling across surveys and identify 10,922 unique tree IDs. Next, we assigned two spatial classifications to these tree IDs: (a) planting configuration, describing planting composition based on neighbours, and (b) planting position, their location within the riparian margin. We then linked these classifications to the long-term survival and height records.

##### (a) Planting configuration

We reconstructed planting configuration separately for each planting line using its earliest available survey, when the spatial arrangement of trees along that line was first recorded. Within this initial record of each line, we identified the species of each focal tree and its immediately adjacent planting positions in the four cardinal directions to determine its original neighbourhood configuration. We then assigned each focal tree a permanent planting-configuration classification linked to its unique tree ID. This allowed us to retain the original configuration across subsequent surveys, even when neighbouring trees were not resampled or were no longer recorded.

For each focal seedling, we identified immediate adjacent planting in each of the four cardinal directions within the planting grid. Focal seedlings were classified as Conspecific when all four neighbouring positions were present (not a border tree) and contained the same species as the focal seedling. They were classified as Mixed when all four neighbours were from a different species to the focal species, were all different from one another, and were all present (not a border tree). This identified 236 Conspecific and 663 Mixed focal seedlings, giving 899 unique focal trees available for comparison.

Monitoring effort varied across surveys because COVID-19 restrictions and subsequent fieldwork changes meant some surveys covered only one riverbank, part of a river margin, or one of the two river sites. Consequently, Mixed and Conspecific planting configurations were not monitored equally throughout the complete survey period, with Conspecific positions rarely resurveyed after survey 5 at PLPE and survey 6 at KNDE. Comparisons between Mixed and Conspecific configurations were therefore restricted to periods of comparable monitoring: surveys 1–5 at PLPE and surveys 1–6 at KNDE.

##### (b) Planting position

We classified planting position independently of planting configuration. Because planting-line lengths varied with the river’s irregular shape, we calculated each tree’s relative location along its planting line and divided the line into three approximately equal sections: River, Middle, and Edge. Riverbank orientation was accounted for so that River consistently represented the section closest to the river and Edge the outermost section of the restored margin.

##### Survival

For survival analyses, we ordered observations chronologically within each tree ID. We defined mortality as the first observed transition from alive to dead. We excluded records after this event, so each seedling contributed only one mortality event and was removed from the risk set after death. If a seedling was not recorded during subsequent surveys but had no observed transition to dead, its disappearance was not classified as mortality, and the seedling was instead treated as no longer observed. After these steps, the planting-configuration survival analysis included 3,683 observations, while the planting-position survival analysis included 44,899 observations.

##### Height (cm)

We first screened height measurements for transcription and unit-entry errors. We corrected 758 measurements identified as recording errors, including inaccurate conversions between metres and centimetres. We then screened longitudinal height records for inconsistencies between successive observations. A measurement was flagged when height decreased by more than 5 cm relative to the preceding observation or exceeded 30,000 cm. For an affected tree ID, observations preceding the first flagged measurement were retained, whereas the flagged measurement and subsequent height records were excluded. Height analyses were restricted to valid measurements from seedlings recorded as alive. After these steps, the planting- configuration height analysis included 2,623 observations, and the planting-position height analysis included 32,230 observations.

### Ecosystem functions associated with enrichment planting

In addition to the long-term dataset described above, we conducted targeted field sampling from January to March 2024 to determine whether ecosystem functions varied with the planting configuration and spatial position. Five to six years after restoration, we did not have enough surviving trees to construct a balanced design across all six originally planted native species. We therefore focused on three species for which we could replicate both Mixed and Conspecific focal trees: *Falcataria moluccana*, *Alstonia scholaris*, and *Peronema canescens*. For each species, we selected three replicate focal trees from each planting configuration within each restoration treatment and river site. This produced 72 focal trees in total (3 replicates × 2 planting configurations × 3 species × 2 treatments × 2 river sites), comprising 36 Mixed and 36 Conspecific trees.

We quantified position within the margin differently for the ecosystem-function measurements than for the long-term dataset, as we were able to assess exact tree location in the margin more accurately in the field. We recorded the GPS location of each selected focal tree and quantified its position continuously as “distance from the river edge (m)”. This provided a direct continuous measure of the spatial gradient across the riparian margin for testing relationships between ecosystem functions and distance from the river.

### Ecosystem functions

#### Tree identification for Planting type and distance from the river

We measured each focal tree, recording predation, leaf herbivory, leaf-litter biomass, total arthropod abundance, and abundance of key herbivore and predator groups (caterpillars, ants, and spiders). We also measured soil chemical properties (cation exchange capacity (CEC), organic carbon, total nitrogen, phosphorus, potassium, and pH) and soil physical properties (bulk density, aggregate stability, friable soil content, and A-horizon depth).

#### Predation

The day before sampling, we collected oil palm fronds and cut them into sections about 20 cm long, each containing six leaflets (three leaflets per side). On the morning of each survey, we attached six live mealworms (*Tenebrio sp*.) of similar size to each frond section using non-toxic adhesive. In the field, we tied one prepared palm frond approximately 1.5 m above ground to the central tree. After 24 hours, the frond was revisited, and the mealworm’s removal rate was recorded as: (i) fully removed, (ii) partially removed, or (iii) not removed.

##### Data processing

Predation was measured as the proportion of each mealworm removed, with fully removed as 1, partially removed as 0.5, and not removed as 0. This then provided a beta-ordinal removal score between 0 and 1.

### Arthropod and herbivory sampling

We conducted herbivory and arthropod community surveys concurrently. We cut one accessible branch with a minimum target length of 3 m and immediately placed it on a white survey sheet (Figure 1).

**Figure 1.**
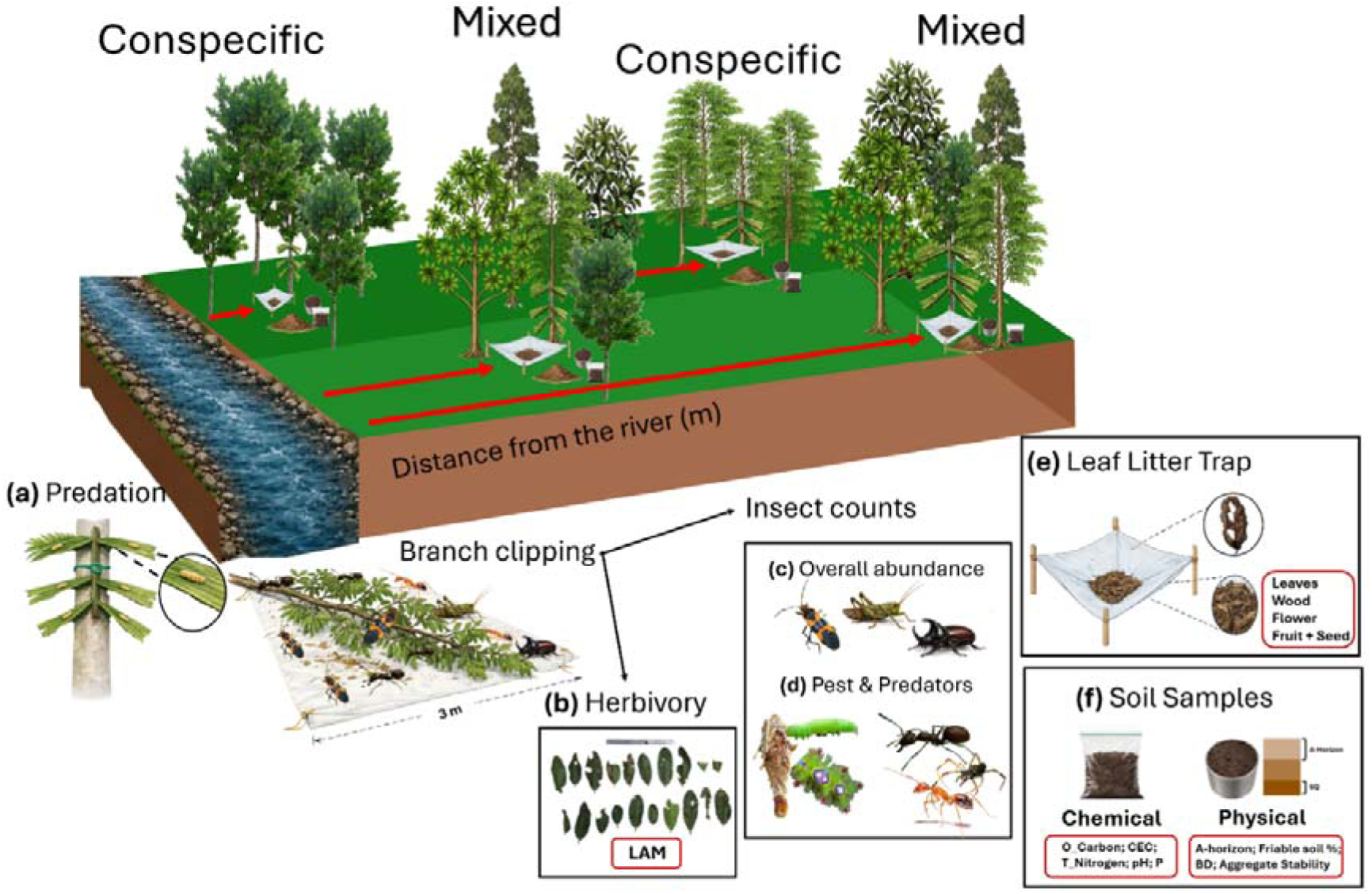
Experimental design and ecosystem-functions measured across two actively restored riparian margins. Active restoration comprised enrichment planting of six native tree species: *Peronema canescens*, *Shorea leprosula*, *Alstonia scholaris*, *Artocarpus integer*, *Calophyllum inophyllum*, and *Albizia chinensis*. Trees were planted within two restoration treatments: Mature Palms + Planting (MP + P), in which remnant mature oil palms were retained alongside enrichment plantings, and Planting Only (P Only), in which mature oil palms were removed prior to enrichment planting. Ecosystem functions were surveyed on three focal enrichment-planted tree species (Sungkai, Sengon, Pulai) across Conspecific (neighbouring trees of the same species) and Mixed (neighbouring trees of different species) planting configurations and along a gradient of distance from the edge (m). Measurements taken at each native tree species comprised: **(a)** predation, assessed using mealworms attached to prepared oil palm fronds; **(b)** foliar herbivory, quantified from 20 leaves per branch-clipping sample using leaf-area measurements in ImageJ; **(c)** total arthropod abundance counted from branch-clipping samples; **(d)** abundance of focal arthropod groups, including the oil palm pest caterpillars *Pteroma pendula* and *Darna pallivitta*, and predators comprising *Oecophylla smaragdina*, other ants, and spiders; **(e)** leaf-litter biomass collected in litter traps over 10 days and separated into leaves, woody material, flowers, reproductive material (seeds and fruit), and herbivory-associated material; and **(f)** soil properties at each location, comprising chemical properties (organic carbon, total nitrogen, available phosphorus, and pH) and physical properties (bulk density, friable-soil proportion, aggregate stability, and A-horizon depth). Figure created by Sacchi Shin-Clayton using Inkscape; copyright- free images sourced from Shutterstock.

All visible arthropods present on the branch were recorded during a timed survey of up to 10 minutes. We identified specimens to order level in the field (Figure 1c). We conducted an additional survey count for key predator and pest taxa commonly found in oil palm plantations. Predator groups recorded included total spider abundance, total ant abundance, and weaver ant (*Oecophylla smaragdina*) abundance. Similarly, we counted caterpillar abundance, split into general caterpillars, identified as bagworms (*Pteroma pendula*), and nettle caterpillars (*Darna pallivitta)*.

After the ten-minute survey, we photographed the entire branch with a scale ruler. We then randomly selected a subset of 20 leaves, placed them on a whiteboard, and photographed them individually, including a 30 cm ruler for scale. Back in the laboratory, we analysed the digital images using ImageJ software (Schneider et al., 2012) to quantify the leaf area loss per leaflet.

### Leaf Litter

We placed a litter trap (1 × 1 m, ∼50 cm above ground) beneath the central tree and within 1 m of the base. We left litter traps in the field for 10 days, then sorted the contents in the laboratory into five categories: leaves, flowers, fruit, seeds, and woody material. We then analysed the contents using two approaches: biomass (weight) and herbivory damage. We measured biomass by weighing the fresh (wet) mass of each category on a precision balance (METTLER TOLEDO©), oven-drying samples (50 °C) to constant mass, and reweighing them to calculate dry biomass. We measured herbivory damage by sorting leaves from each trap into either damaged or untouched categories. We then calculated the proportion of damage as damage/total number of leaves. For Sengon, we based herbivory assessments on a random subsample of 100 leaves per trap, given the high abundance of leaflets in litter samples.

### Soil samples

#### In field measurements

At the base of the central tree (< 50 cm distance), a soil core was collected using a cylindrical corer (height 4.1cm x 7.3 cm width = 171.7 cm^3^). We placed the pipe vertically on the soil surface and gently drove it into the ground with a mallet until the upper edge was flush with the soil surface. We then extracted the complete core carefully using a trowel and transferred the contents to the lab in a labelled Ziploc bag.

We completed a rapid Visual Evaluation of Soil Structure (VESS) assessment in the field, as described by Batey et al. (2015). 50 cm east of the soil core samples, we initially excavated a pit approximately 40 cm deep, 35 cm wide, and 30 cm long. The depth of the topsoil (hereafter A-horizon) was measured from the soil surface (Figure 1f). Adjacent to the pit, we cut and removed a soil block (∼25 cm deep, 20 cm wide, and 15 cm long) and placed it on a plastic tray for VESS analysis (Figure 1f). We identified soil layers, measured the depth of each identified layer, and assigned each layer a soil quality (Sq) score from 1 to 5 using the standard VESS framework. We calculated friable soil content as the cumulative thickness of layers assigned a VESS score of 1 (representing the most friable, well-structured soil), then summed it and expressed it as a percentage of the total profile depth (25 cm). Finally, we collected a composite sample (∼1 kg) of all the layers into a Ziploc bag and transported it back to the lab.

#### Soil Laboratory protocol

##### Bulk density

Immediately after returning from the field, we weighed the wet mass of each soil core sample (METTLER TOLEDO©). On the same day, we oven-dried the samples (Memmert© incubator) at 105°C until the weight was constant, and then recorded the final dry weight to determine bulk density (BD) using the standard calculation (BD = (wet weight – dry weight)/171.7 cm^3^).

##### Soil nutrients

The composite soil samples taken from the field were air-dried until moisture was removed (∼14 days). The contents were sieved through an ISO 3310-1/ASTM E.11 sieve (ø 200 mm / 203 mm (8 inch)) and a 125 mm-20 µm mesh filter (Cole-Parmer©). Samples were sent to the Sinar Mas Analytical Laboratory, Bogor, Indonesia, for analysis of soil physicochemical properties. Soil water content was measured using the gravimetric method with MA45 Moisture Analyser Sartorius Type(FAO, 2023b). Texture was determined using the hydrometer method (Bouyoucos, 1962); pH was measured potentiometrically; organic carbon by Walkley–Black titration (Walkley & Black, 1934); soil aggregate % (Eijkelkamp, 2022) and total nitrogen by the Kjeldahl method (Bremner, 1960). Reserve P, K and Mg were determined following extraction with 25% HCl; available P using Bray I extraction (Bray & Kurtz, 1945) and cation exchange capacity (CEC) and exchangeable base cations (Ca, Mg, K and Na) using ammonium acetate (NH OAc) at pH 7 (Motsara & Roy, 2008). Exchangeable H and Al were determined by titration (Motsara & Roy, 2008).

##### Data processing

We applied a standardised multivariate reduction workflow to the full soil chemical dataset (16 variables) to identify key variables determining soil characteristics and select a minimum data set (MDS) for subsequent analyses (Andrews et al., 2002; Shokr et al., 2025). We first performed a Principal component analysis (PCA) to visualise general trends, then systematically excluded components with eigenvalues 1 until approximately 70–80% of the total variance was explained (PC1 – PC4 = 73.4% of the variance) (Andrews et al., 2002; Kaiser, 1960). Variables were evaluated using factor loadings (≥0.5), percentage contribution and squared cosine values (cos²) (Andrews et al., 2002; Cantú et al., 2007). We then reduced highly correlated variables (|r| ≥0.7) based on ecological relevance and redundancy to produce the final MDS (Li et al., 2024), comprising organic carbon, pH, CEC, exchangeable available phosphorus, and total nitrogen (Supplementary Table S7).

### Statistical analysis

All statistical analyses were performed in RStudio version 4.4.2 (Posit team, 2023).

#### Seedling survival and height

We assessed seedling survival using discrete-time survival analysis. Each survey in which a seedling remained under observation represented a Bernoulli trial, with observations while the seedling remained alive coded as 0, and the first observed mortality event coded as 1; observations following mortality were excluded. Seedlings that were no longer recorded in subsequent surveys without an observed mortality event were treated as no longer observed rather than assumed to have died. Mortality was analysed using binomial generalized linear mixed-effects models (GLMMs) with a logit link, fitted using the ‘glmmTMB’ package (Brooks et al., 2017). For analyses of planting configuration, comparisons between Mixed and Conspecific plantings were restricted to surveys with comparable monitoring effort (surveys 1–5 at PLPE and 1–6 at KNDE). Models included restoration treatment, planting configuration or planting position, survey period, and supported interactions among these predictors as fixed effects, with site included as an additional fixed effect and planting plot (‘Plot_ID’) as a random intercept.

We analysed seedling height using linear mixed-effects models, with log-transformed height as the response variable. Models included restoration treatment, planting configuration or planting position, survey period, and their interactions as fixed effects, with seedling identity included as a random intercept to account for repeated height measurements of the same individual. Planting-configuration height analyses were restricted to the same periods of comparable Mixed and Conspecific monitoring used for the survival analysis.

#### Ecosystem functions

Ecosystem functions for predation, leaf litter biomass, arthropod abundance, and soil variables were modelled using general linear mixed models (GLMMs) (Brooks et al., 2017). Each ecosystem survey was the response variable, with restoration treatment and planting type as fixed effects with an interaction, and river site: tree species nested as a random effect. For distance from the river, we again used GLMMs, with restoration treatment and distance as fixed effects and their interaction.

We modelled herbivore predation and herbivory on branch clippings as proportions (0–1) and analysed them using GLMMs with an ordered beta distribution and a logit link function. For herbivory, leaf area loss was averaged across 20 leaf clippings sampled per branch, and Plant ID was included as a random effect to account for repeated measurements from the same plant. We analysed predator and pest species abundance counts using negative binomial models because the dataset had many zeros, but with the same model structure: abundance ∼ Treatment * Planting type * Distance. We modelled bagworm and nettle caterpillar abundance without interactions among the fixed effects due to singularity issues, instead including additive effects of treatment, planting type, and distance.

## Results

Seedling survival did not differ between the planting types (GLMM: χ*²* = 0.024, *df* = 1, *p* = 0.878; Figure 2A), or in its interaction with either treatment (χ*²* = 0.024, *df* = 1, *p* = 0.877) or survey (χ*²* = 1.3, *df* = 1, *p* = 0.935). However, position within the margin interacted significantly with restoration treatment (χ*²* = 6.413, *df* = 2, *p* = 0.041), but not with survey (χ*²* = 6.413, *df* = 16, *p* = 0.041; Figure 2b).

**Figure 2.**
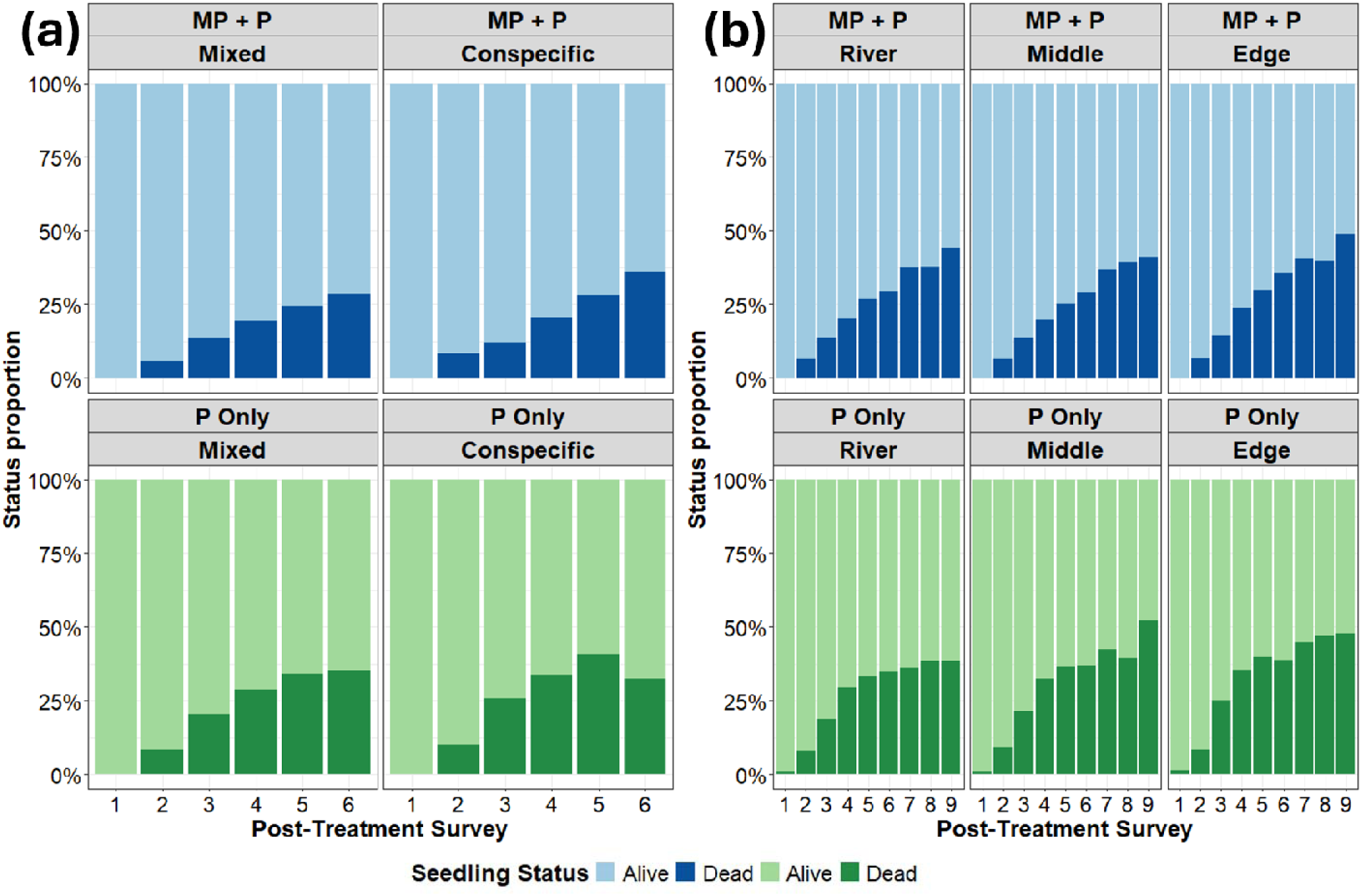
Status (Alive or Dead) of the plantings in the two actively restored treatments: Mature Palms and Planting (top) and Planting Only (bottom). Enrichment plantings included an equal number of the six native tree species *Peronema canescens*; *Shorea leprosula*; *Alstonia scholaris*; *Artocarpus integer*; *Calophyllum inophyllum*; *Albizia chinensis* between consecutive post-treatment surveys (surveys 1–6 following enrichment planting for planting type and surveys 1 - 9 for position). (a) Tree status grouped by planting configuration (Conspecific: same species planted around the focal tree, Mixed: different species as neighbours). **(b)** Tree status grouped by their position within the riparian margin (river, middle, edge closest to the oil palm plantation).

**Figure 3.**
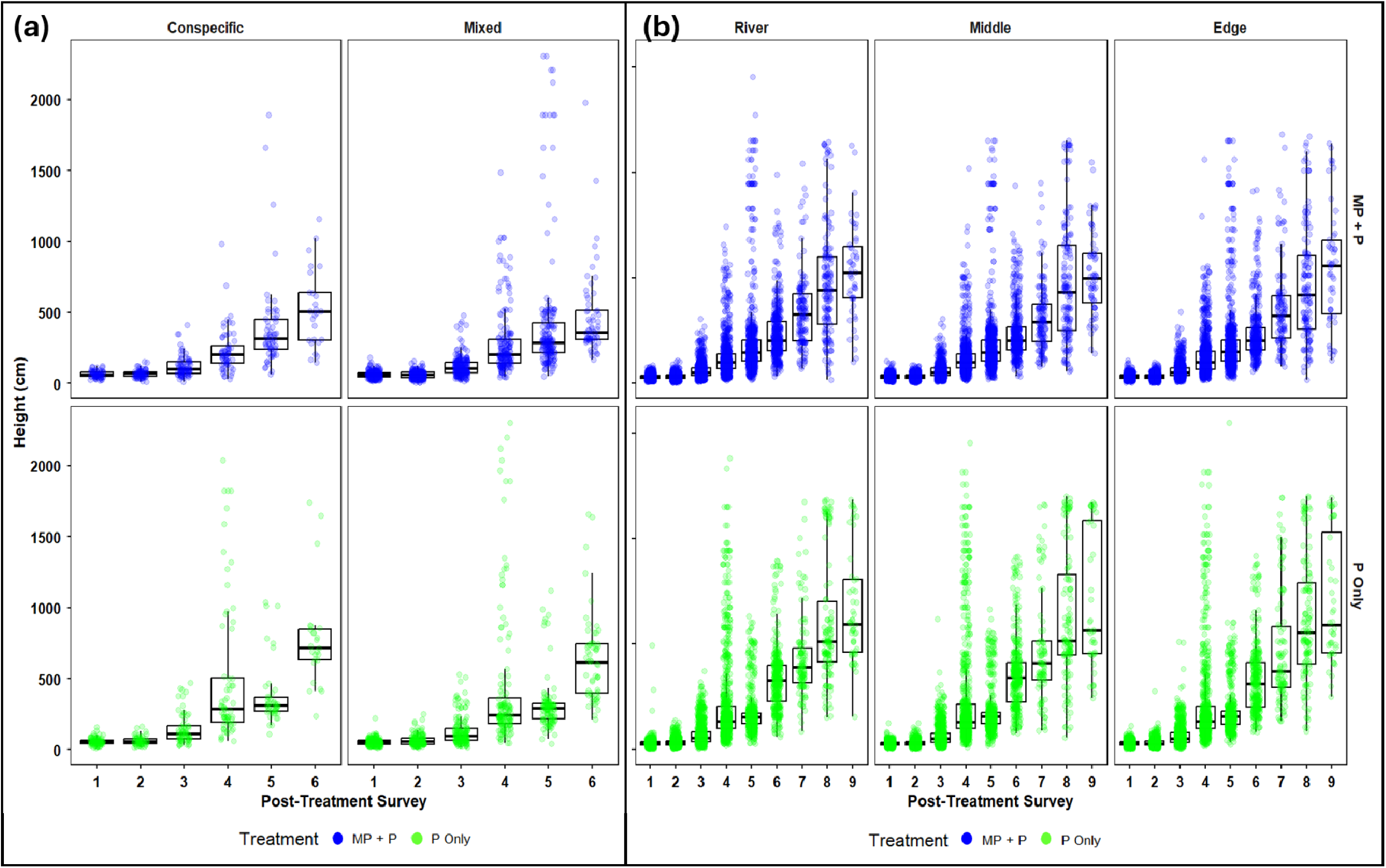
Seedling height (cm) across actively restored riparian buffers. Differences between enrichment plantings established beneath retained mature oil palms (MP + P, top plots) and enrichment plantings with mature oil palms removed from the margin (P Only, bottom plots) for each of the six native tree species *Peronema canescens*; *Shorea leprosula*; *Alstonia scholaris*; *Artocarpus integer*; *Calophyllum inophyllum*; *Albizia chinensis* between consecutive survey periods (surveys 1–6 following enrichment planting for planting type and surveys 1 - 9 for position). **(a)** The enrichment plantings are grouped by planting configuration (Conspecific: same species planted around the focal tree, Mixed: different species planted around the focal tree). **(b)** Seedling height was grouped by their position within the riparian margin (river, middle, edge).

**Figure 4.**
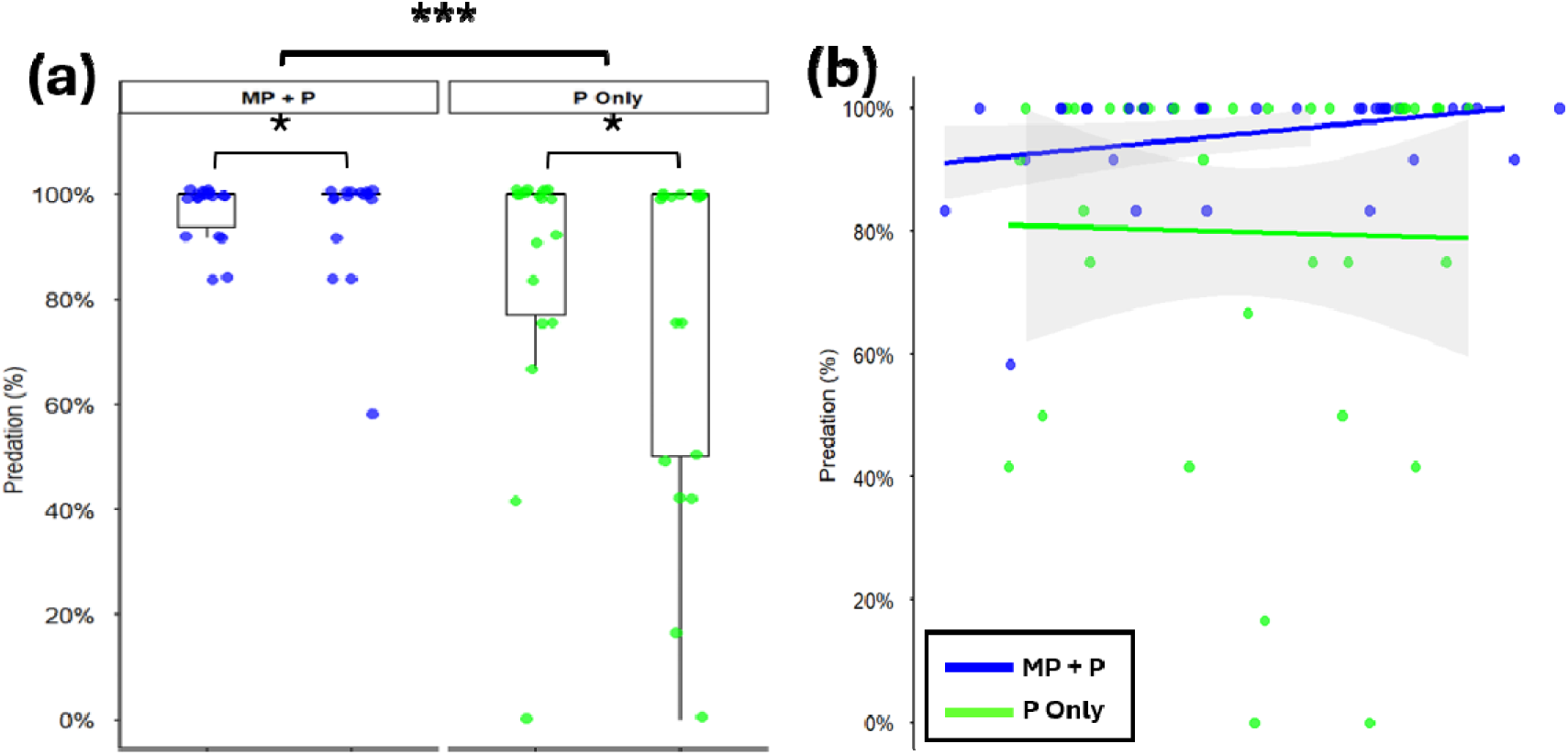
Mealworm predation rate quantified as mealworm removal (0–6; intact to completely removed). **(a)** Predation rate is grouped by restoration treatment (MP + P: Mature Palms and Plantings, P Only: Plantings Only) and planting configuration (Conspecific: same species planted around the focal tree, Mixed: different species as neighbours). **(b)** Predation rate is plotted by distance (m) to the river, and lines represent restoration treatments (Blue = MP + P and Green = P Only), with 95% confidence intervals shown by grey shading.

**Figure 5.**
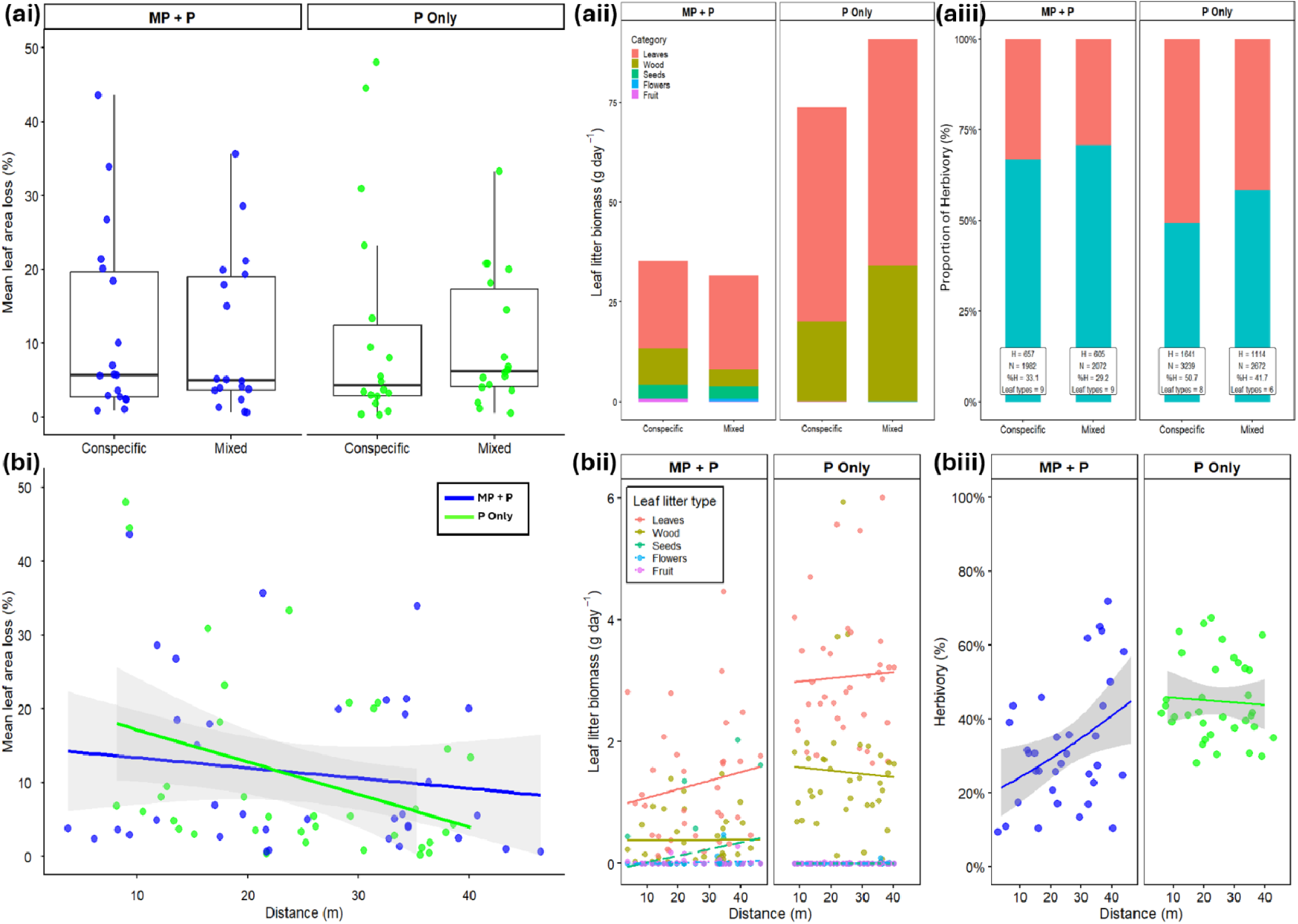
Effects of restoration treatment, planting configuration, and distance from the river on herbivory and leaf-litter biomass. (ai) Mean leaf area loss (%) and (aii) mean daily leaf-litter biomass by litter category for conspecific and mixed- species plantings. (aiii) Proportion of leaves with and without herbivory; insets show total leaves (*N*), leaves with (*H*) and without (*NH*) herbivory, and mean leaves sampled per plant. (bi–biii) Relationships between distance from the river and (bi) mean leaf area loss, (bii) biomass of individual litter categories, and (biii) percentage of leaves with herbivory. Points in (ai) represent plant means; boxplots show medians, interquartile ranges, and whiskers. Lines in (bi–biii) show fitted linear relationships, with shaded areas indicating 95% confidence intervals where shown. MP + P = enrichment planting with mature oil palms retained; P Only = enrichment planting with mature oil palms removed.

**Figure 6.**
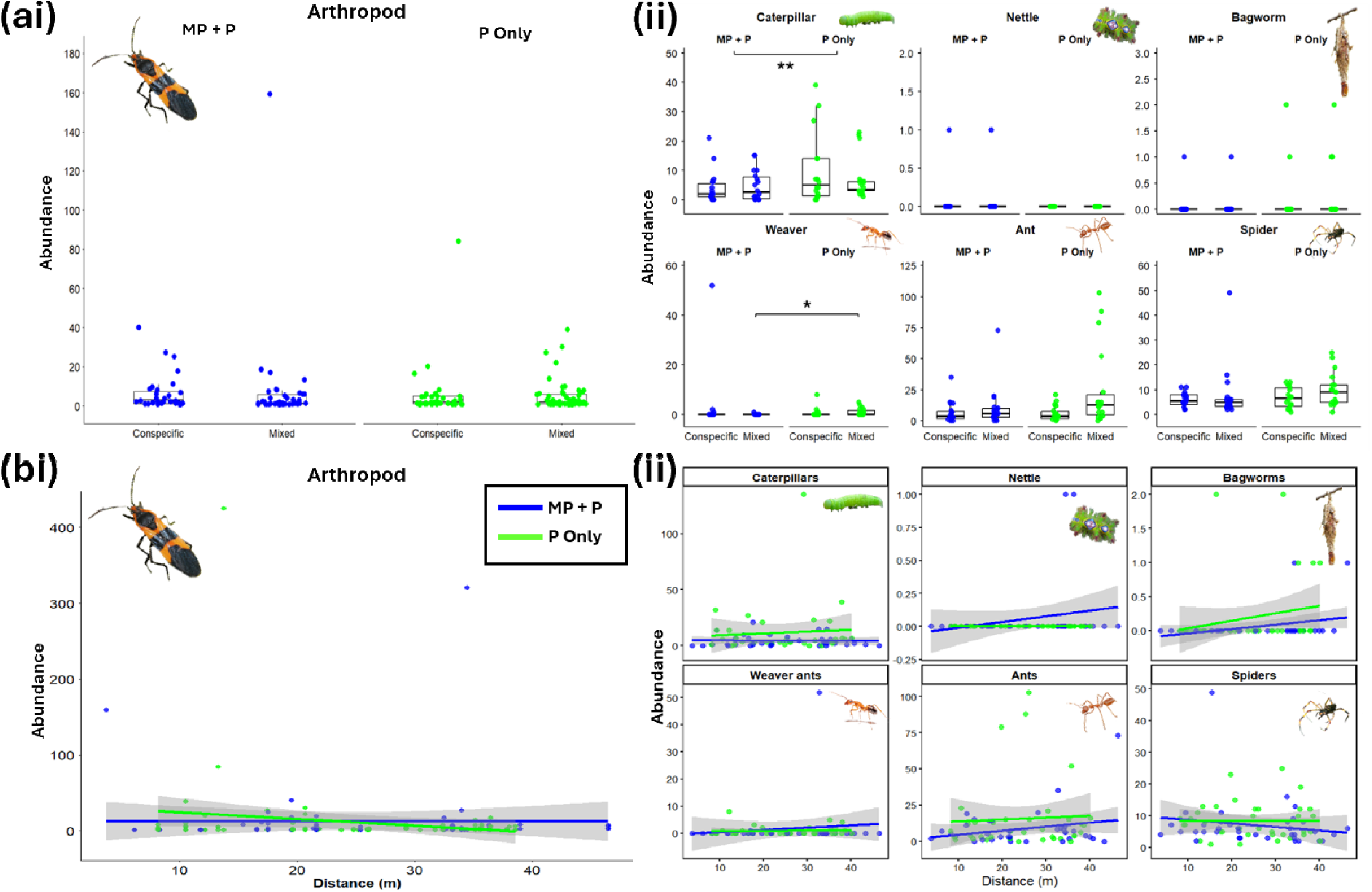
Arthropod abundance grouped by planting type and position within the riparian margins. The top row **(a)** shows abundances in relation to planting configuration (Conspecific and Mixed), while the bottom row **(b)** shows abundances in relation to distance from the river (m). (**ai** and **aii)** Abundance is grouped by restoration treatment (MP + P: Mature Palms and Plantings, P Only: Plantings Only) and planting configuration (Conspecific: same species planted around the focal tree, Mixed: different species as neighbours). (**bi** and **bii)** Abundance is plotted by distance in metres to the river, and lines are restoration treatments (Blue = MP + P and Green = P Only), with 95% confidence intervals shown by grey shading.

**Figure 7.**
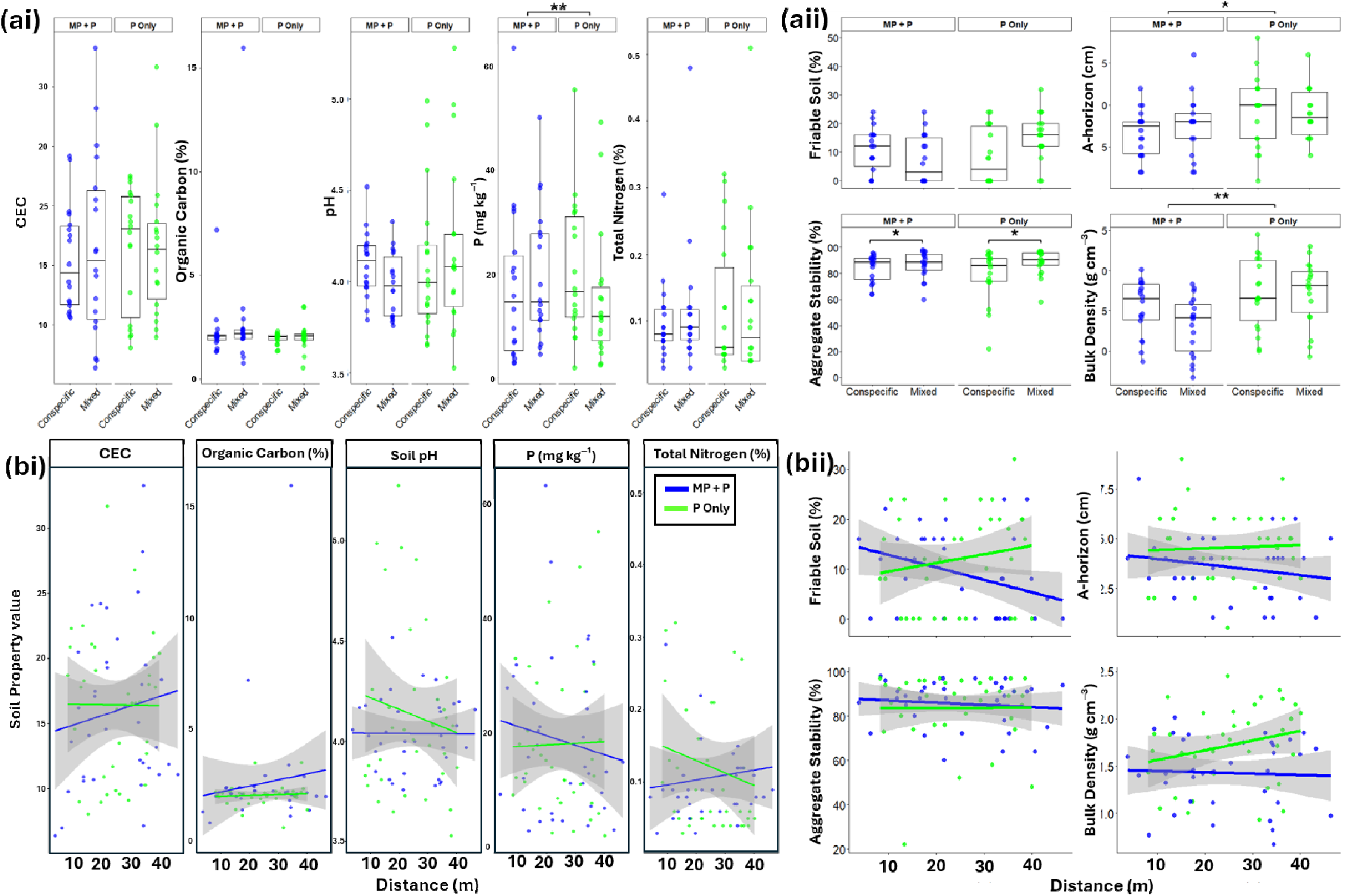
Soil chemical and physical properties measured at the base of enrichment-planted trees within actively restored riparian margins. The top row **(a)** shows soil properties in relation to planting configuration (Conspecific and Mixed), while the bottom row **(b)** shows soil properties in relation to distance from the river (m). **(ai** and **bi)** Soil chemical properties, including cation exchange capacity (CEC), soil organic carbon (%), pH, phosphorus concentration (mg kg^⁻¹^), and total soil nitrogen (%). **(aii** and **bii)** Soil physical properties, including the proportion of the soil profile classified as friable using the Visual Evaluation of Soil Structure (VESS) protocol from a 25-cm-deep extraction block; A-horizon depth (cm), measured as the depth of the topsoil layer within the 0–25 cm extraction block; aggregate stability (%), measured from soil samples collected within each buffer core plot; and bulk density (g cm^⁻³^).

Seedling position within the margin had a greater impact on survival in the Plantings Only margin than in Mature Palms and Planting, which did not have different survival rates (River, Middle, Edge, all p > 0.05). Within the Planting Only treatment, survival decreased from the river towards the edge (River: 5.74%; Middle: 6.74%; Edge: 7.40%). Therefore, survival was higher at the River than at the Edge (OR = 0.763 ± 0.07 SE, *p* = 0.0084), whereas the remaining comparisons were not significantly different (River vs Middle: OR = 0.842 ± 0.081 SE, p = 0.173; Middle vs Edge: OR = 0.905 ± 0.083, *p* = 0.524).

Treatment effects varied between survey periods (χ*²* = 143.607, *df* = 8, *p* < 0.0001; Figure 2b), with survival being significantly higher in the Mature Palms and Planting margin than in Planting Only during survey 3 (7.57% vs 15.60%; OR = 0.443, p < 0.0001), survey 4 (8.75% vs 14.32%; OR = 0.574, p < 0.0001), and survey 8 (14.02% vs 21.00%; OR = 0.613, p = 0.0014). In contrast, survival was lower in Mature Palms and Planting during survey 5 (8.76% vs 6.41%; OR = 1.40, *p* = 0.0007), but not significant for the other survey intervals (all p > 0.05). Planting position also differed significantly between survey periods, with survival higher at the river than at the edge during surveys 3 (OR = 0.749, *p* = 0.0009), 4 (OR = 0.771, *p* = 0.0104), and 6 (OR = 0.564, *p* = 0.0025). Survival was also higher in the Middle compared to the Edge for survey 4 (OR = 0.808, *p* = 0.0475), and the remaining comparisons were not significant (all p > 0.05).

Seedling height did not differ by planting configuration (χ*²* = 1.81, *df* = 1, *p* = 0.179), and planting configuration did not interact with either treatment (χ*²* = 1.01, *df* = 1, *p* = 0.316) or survey period (χ*²* = 3.26, *df* = 5, *p* = 0.660). In contrast, seedling height varied significantly with the interaction between restoration treatment and survey period (χ*²* = 154.87, *df* = 5, *p* < 0.001). At survey 1, seedlings were taller in Mature Palm and Planting than in Planting Only (*OR* = 1.218, *p* < 0.001), whereas by surveys 4 and 6 seedlings were taller in Planting Only (*OR* = 0.663 and 0.641, respectively; both *p* < 0.001).

Seedling height did not differ significantly between treatments at surveys 2, 3, or 5 (all p > 0.05). Seedling height varied significantly with the interaction between planting position and survey period (χ*²* = 27.66, *df* = 16, *p* = 0.035). Post-hoc comparisons showed that at survey 2, seedlings at the River were significantly taller than those at both Middle (58.1 vs 55.2 cm; *OR* = 1.054, *p* = 0.006) and Edge (58.1 vs 55.7 cm; *OR* = 1.043, *p* = 0.027), whereas Middle and Edge did not differ (*p* = 0.816). No significant differences among planting positions were detected during the other survey periods (all *p* > 0.05). Lastly, there was no significant interaction between treatment and position (χ*²* = 0.84, *df* = 2, *p* = 0.659) or overall effect of planting position (χ*²* = 2.38, *df* = 2, *p* = 0.304).

### Ecosystem functions

#### Predation

Mealworm removal rate by planting type differed significantly with distance from the river (χ*²* = 5.339, *df* = 1, *p* = 0.021). Post hoc slope analyses showed that mealworm removal decreased significantly with increasing distance from the river for conspecific plantings under Planting Only (slope = −0.051 ± 0.024, *z* = −2.118, *p* = 0.034), whereas no significant relationship with distance was detected for conspecific plantings under Mature Palms and Planting (slope = 0.009 ± 0.021, *p* = 0.661) or mixed plantings under either Mature Palms and Planting (slope = 0.032 ± 0.027, *p* = 0.232) or P Only (slope = 0.033 ± 0.020, *p* = 0.092). Within Planting Only, the distance slope was significantly more negative for conspecific than mixed plantings (difference in slopes = −0.084 ± 0.033, *z* = −2.589, *p* = 0.010), whereas their slopes did not differ under Mature Palms and Planting (difference = −0.023 ± 0.034, *z* = −0.680, *p* = 0.497).

Overall, estimated mealworm removal differed between treatments (GLMM: χ*²* = 20.950, *df* = 1, *p* < 0.0001) and was higher under Mature Palms and Planting (Mature Palms and Planting; 85.3%, 95% CI = 76.6–91.2%) than under Planting Only (P Only; 62.6%, 95% CI = 53.2–71.1%), corresponding to 3.47-times greater odds of removal under Mature Palms and Planting (OR = 3.47 ± 0.986, *z* = 4.381, *p* < 0.0001).

Planting type affected mealworm removal rates (χ*²* = 9.549, *df* = 1, *p* = 0.002); however, the marginal contrast averaged across treatments and distance did not detect a significant difference between conspecific and mixed plantings (OR = 1.60 ± 0.530, *z* = 1.416, *p* = 0.157), consistent with planting type varying with distance. Finally, there was no overall significant effect of distance (χ*²* = 0.342, *df* = 1, *p* = 0.559), treatment × planting type (χ*²* = 3.038, *df* = 1, *p* = 0.081), treatment × distance (χ*²* = 1.822, *df* = 1, *p* = 0.177), or treatment × planting type × distance interactions (χ*²* = 1.746, *df* = 1, *p* = 0.186).

#### Herbivory

For branch clippings, herbivory damage did not differ between restoration treatments (χ*²* = 0.025, *df* = 1, *p* = 0.875) or planting types (χ² = 0.162, *df* = 1, *p* = 0.687), and there were no significant interactions (all *p* ≥ 0.102). However, herbivory decreased significantly with increasing distance from the river (χ*²* = 5.034, *df* = 1, *p* = 0.025; slope = −0.027 ± 0.012 SE, *z* = −2.377, *p* = 0.018), with no evidence that this relationship differed between restoration treatments (Treatment × Distance: χ*²* = 2.078, *df* = 1, *p* = 0.149; Figure 5bi).

For leaf litter contents, herbivory damage and biomass differed between the restoration treatments (Herbivory: *X^2^*= 16.677, *df* = 1, *p <* 0.0001; Biomass: *X^2^* = 73.06, *df* = 1, *p <* 0.0001), but not between the planting types (Herbivory: *X^2^* = 1.196, *df* = 1, *p =* 0.274; Biomass: *X^2^ =* 1.236, *df* = 1, *p =* 0.266), distance (Herbivory: *X^2^*= 2.219, *df* = 1, *p =* 0.136; Biomass: *X^2^* = 0.622, *df* = 1, *p =* 0.43), or the interactions between treatment x planting (Herbivory: *X^2^* = 0.684, *df* = 1, *p =* 0.408; Biomass: *X^2^* = 1.099, *df* = 1, *p =* 0.294), treatment x distance (Herbivory: *X^2^* = 3.05, *df* = 1, *p =* 0.081; Biomass: *X^2^* = 1.908, *df* = 1, *p =* 0.167), planting x distance (Herbivory: *X^2^* = 0.534, *df* = 1, *p =* 0.465; Biomass: *X^2^* = 2.074, *df* = 1, *p =* 0.15), or treatment x planting x distance (Herbivory: *X^2^* = 1.828, *df* = 1, *p =* 0.176; Biomass: X^2^ = 0.259, *df* = 1, *p =* 0.611). Between the treatments, herbivory and biomass and were higher in the Planting Only margin (Herbivory: estimate = -0.566 ± 0.135, t = -4.195, *p <* 0.0001; Biomass: *OR =* 0.379 ± 0.043, *z =* -8.638, *p <* 0.0001). Litter type also varied between treatments, with all five categories having higher biomass in Planting Only margins than in Mature Palms and Planting (all *p <* 0.0001).

#### Arthropod abundance

Total arthropod abundance was influenced significantly by the interaction between planting type and distance from the river (GLMM: χ*²* = 12.410, *df* = 1, *p* < 0.0001). Post hoc slope comparisons supported the planting type × distance interaction: under both restoration treatments, the relationship between arthropod abundance and distance was significantly more negative for conspecific than mixed plantings (Mature Palms and Planting: difference in slopes = −0.108 ± 0.043, *z* = −2.52, *p* = 0.012; Planting Only: difference in slopes = −0.112 ± 0.047, *z* = −2.38, *p* = 0.017). Thus, arthropod abundance declined more strongly with increasing distance from the river in conspecific than mixed plantings, irrespective of restoration treatment. In contrast, there were no significant overall effects of restoration treatment (χ*²* = 0.678, *df* = 1, *p* = 0.411), planting type (χ*²* = 2.071, *df* = 1, *p* = 0.150), or distance (χ*²* = 3.235, *df* = 1, *p* = 0.072). All remaining interactions were also not significant (all *p* > 0.05)

#### Predator and Pest groups

Responses of individual arthropod groups varied. Overall caterpillar abundance exhibited a significant three-way interaction between restoration treatment, planting type, and distance (χ*²* = 5.678, *df* = 1, *p* = 0.017). Within Mature Palm and Planting, the relationship between caterpillar abundance and distance differed significantly between Mixed and Conspecific plantings (difference in slopes = −0.099 ± 0.038, *p* = 0.009). Caterpillar abundance increased with distance from the river in Mixed plantings (slope = 0.074 ± 0.032, *p* = 0.020), but did not change with distance in Conspecific plantings (slope = −0.025 ± 0.025, *p* = 0.322). In Planting Only, distance slopes did not differ between Mixed and Conspecific plantings (*p* = 0.387), and neither slope differed significantly from zero (both *p* > 0.17). Caterpillar abundance differed between treatments (χ*²* = 11.659, *df* = 1, *p* = 0.0006) and was higher in the Planting Only margin (*response* = 8.98 ± 3.09 SE) than in the Mature Palm and Planting (*response* =3.42 ± 1.21 SE; OR = 0.381, *p* < 0.001). All the remaining interactions and single factors were not significant (all > 0.05).

General ant abundance, however, differed significantly between planting types (χ*²* = 12.288, *df* = 1, *p* = 0.0004) and showed an overall effect of distance (χ*²* = 3.907, *df* = 1, *p* = 0.048). Despite the significant overall effect of distance, the estimated marginal trend did not indicate a significant change in ant abundance with increasing distance from the river (slope = 0.015 ± 0.012, *z* = 1.20, *p* = 0.229). Between restoration treatments (χ*²* = 2.054, *df* = 1, *p* = 0.152), and there were no significant interactions (all *p* > 0.05).

Bagworm and nettle caterpillars, spiders, and weaver ants generally showed no significant responses to restoration treatment, planting type, or distance from the river. Bagworm and nettle caterpillar abundance did not vary with planting type (Bagworm: χ*²* = 0.006, *df* = 1, *p* = 0.937; Nettle: χ² = 0.031, *df* = 1, p = 0.860) or distance (Bagworm: χ*²* = 0.027, *df* = 1, *p* = 0.870; Nettle: χ*²* = 1.377, *df* = 1, *p* = 0.241), while spiders and weaver ants showed no significant effects of treatment, planting type, or distance (all *p* > 0.05). No interactions were significant for spiders or weaver ants (all *p* > 0.05).

#### Soil chemical profile

Soil organic carbon was significantly influenced by the interaction between restoration treatment and planting type (GLMM, *t*-family: χ*²* = 12.351, *df* = 1, *p* = 0.0005). This indicates the overall treatment difference did not work in isolation. Within mixed plantings, organic carbon was significantly higher under Mature Palms and Planting (2.23 ± 0.02) than Planting Only (2.04 ± 0.03; *estimate* = 0.186 ± 0.043, *z* = 4.380, *p* < 0.0001), whereas there was no difference between treatments within conspecific plantings (MP + P = 2.08 ± 0.03; P Only = 2.08 ± 0.03; *estimate* = −0.005 ± 0.040, *z* = −0.121, *p* = 0.904). Accordingly, although mixed plantings had numerically higher organic carbon than conspecific plantings when averaged across treatments, this marginal difference was not significant (Conspecific − Mixed: *estimate* = −0.052 ± 0.029, *z* = −1.790, *p* = 0.074).

Organic carbon also showed a significant interaction between planting type × distance, indicating the relationship depended on the planting configuration (χ*²* = 7.421, *df* = 1, *p* = 0.006). Under Mature Palms and Planting, the distance slope was significantly more negative in mixed than conspecific plantings (Conspecific − Mixed: *slope* = 0.0090 ± 0.0032 m^⁻¹^, *z* = 2.807, *p* = 0.005). Specifically, organic carbon declined significantly with distance in mixed plantings (*slope* = −0.0111 ± 0.0023 m^⁻¹,^ *z* = −4.804, *p* < 0.0001), whereas no significant relationship with distance was detected in conspecific plantings (*slope* = −0.0021 ± 0.0022 m^⁻ ¹^, *p* = 0.342). Under Planting Only, the distance slopes did not differ significantly between conspecific and mixed plantings (difference in slopes = 0.0036 ± 0.0047 m^⁻¹^, *z* = 0.769, *p* = 0.442), and neither planting type showed a significant individual relationship with distance (Conspecific: slope = −0.0015 ± 0.0032, *p* = 0.650; Mixed: slope = −0.0051 ± 0.0033, *p* = 0.128). Carbon also declined overall with increasing distance from the river (*slope* = −0.0049 ± 0.0014 m^⁻¹^, *z* = −3.543, *p* = 0.0004).

In contrast, treatment × distance (χ*²* = 1.388, *df* = 1, *p* = 0.239) and treatment × planting type × distance (χ*²* = 0.940, *df* = 1, *p* = 0.332) interactions were not significant. Across planting types, organic carbon was higher under Mature Palms and Planting (2.15 ± 0.02) than Planting Only (2.06 ± 0.02; MP + P − P Only: *estimate* = 0.091 ± 0.029, *z* = 3.133, *p* = 0.002).

#### Soil physical metrics

A-horizon depth differed significantly between restoration treatments, with a deeper A-horizon under Planting Only (P Only; 4.49 ± 0.28 cm) than under Mature Palms and Planting (MP + P; 3.58 ± 0.28 cm). The A-horizon was estimated to be 0.91 cm shallower under Mature Palm and Planting than Planting Only (MP + P − P Only: estimate = −0.914 ± 0.397 cm, *t* = −2.299, *df* = 61, *p* = 0.025). No other effects or interactions were significant for A-horizon depth (all *p* > 0.05).

Bulk density exhibited a significant treatment × planting type × distance interaction (χ*²* = 4.00, *df* = 1, *p* = 0.045), indicating that treatment differences in the relationship between bulk density and distance depended on planting type. Bulk density was significantly lower under Mature Palm and Planting than Planting Only (MP + P − P Only: estimate = −0.166 ± 0.049 g cm^⁻³^, *t* = −3.405, *p* = 0.001), with estimated marginal means of 0.811 ± 0.034 and 0.976 ± 0.034 g cm^⁻ ³^, respectively. Within mixed plantings, distance slopes differed significantly between restoration treatments (difference in slopes = −0.0173 ± 0.0069 g cm^⁻³^ m^⁻ ¹^, *t* = −2.52, *p* = 0.014). Under Planting Only, bulk density increased with distance from the river (slope = 0.0116 ± 0.0055 g cm^⁻³^ m^⁻¹^, 95% CI = 0.0005–0.0226), whereas no significant relationship with distance was detected under Mature Palm and Planting (slope = −0.0058 ± 0.0041 g cm^⁻ ³^ m^⁻¹^, 95% CI = −0.0140–0.0024). In contrast, distance slopes did not differ between restoration treatments within conspecific plantings (*p* = 0.836).

Soil aggregate stability differed significantly between planting types (χ*²* = 4.544, *df* = 1, *p* = 0.033), with greater aggregate stability in mixed than conspecific plantings (Conspecific − Mixed: estimate = −6.49 ± 3.10 percentage points, *p* = 0.041). No other effects or interactions were significant for aggregate stability (all *p* > 0.05).

Finally, treatment × planting type (χ*²* = 10.128, *df* = 1, *p* = 0.002) and treatment × distance interactions (χ*²* = 5.083, *df* = 1, *p* = 0.024) significantly influenced the proportion of friable soil, while no remaining effects or interactions were significant (all *p* > 0.05). Post hoc comparisons showed that treatment effect depended on planting type. Within mixed plantings, friable soil was more than twice as abundant under Planting Only (15.26 ± 1.93%) as under Mature Palm and Planting (6.85 ± 1.91%), a difference of 8.41 percentage points (MP + P − P Only: estimate = −8.41 ± 2.63, *t* = −3.199, *p* = 0.002). In contrast, friable soil did not differ significantly between Mature Palm and Planting (11.63 ± 1.92%) and Planting Only (8.27 ± 1.93%) within conspecific plantings (estimate = 3.37 ± 2.65, *t* = 1.272, *p* = 0.208).

The relationship between friable soil and distance from the river also differed significantly between restoration treatments (difference in slopes, MP + P − P Only = −0.442 ± 0.189, *t* = −2.341, *p* = 0.023). Under Mature Palm and Planting, the proportion of friable soil decreased significantly with increasing distance from the river (slope = −0.288 ± 0.117, 95% CI = −0.522 to −0.055), whereas no significant relationship with distance was detected under Planting Only (slope = 0.153 ± 0.149, 95% CI = −0.145 to 0.452).

## Discussion

Position within the riparian margin and the spatial configuration of enrichment plantings influenced some of the ecosystem functions measured in our study. Seedling mortality and height did not differ between mixed and conspecific plantings, but mixed plantings were associated with greater ant abundance and greater soil aggregate stability. Seedling height also differed by their position in the margin, but only when analysed alongside treatment and time. However, planting configuration did not consistently affect all measured responses; several ecosystem functions were more strongly associated with position within the riparian margin or broader differences among restoration treatments.

### Varied responses to planting configurations

Mortality did not differ between conspecific and mixed-species neighbourhoods, suggesting that local neighbour identity had little influence on survival during the study period. A similar pattern emerged in another enrichment-planting experiment in Sumatran oil palm landscapes, where greater planted-tree diversity increased growth and stand-level productivity but did not affect mortality (Zemp et al., 2019). Together, these results suggest that, within enrichment plantings in oil palm landscapes, planting diversity may be expressed more strongly through tree growth and productivity than through survival, at least within the timeframe of our study.

This lack of effect of planting configuration contrasts with the conspecific negative density dependence (CNDD) reported in natural tropical forests. In these more natural systems, experimental studies provide strong evidence that density dependence influences survival and recruitment. For example, in a study in a Central American rainforest, Bagchi et al. (2010) found rapid density-dependent mortality following germination, with fungal pathogens causing 90–100% mortality within four weeks at high seedling densities. Similarly, Hazelwood et al. (2021) found that fungal pathogens were the strongest contributors to negative density-dependent mortality among tropical tree seedlings they monitored for ten years in Manu National Park, southeast Peru. However, the strength of CNDD varies among species, environmental conditions and life stages (Comita et al., 2014; Zhu et al., 2015). Negative conspecific effects tend to be strongest during early life stages and can weaken as trees develop (Zhu et al., 2015). Our experimental design also differs from many tests of the Janzen–Connell mechanism, as we deliberately planted seedlings at equal spacing as a single cohort, and planting configuration altered the identity of neighbouring trees rather than exposing recruits to gradients of conspecific density or proximity to established reproductive conspecific adults.

Accordingly, the conditions promoting natural-enemy-mediated density dependence may have been less pronounced than in naturally regenerating tropical forests. Our results therefore do not provide evidence for CNDD within these enrichment plantings, but they also do not contradict its importance during other stages or under the higher conspecific densities characteristic of natural recruitment. Thus, the agreement between our results and those of Zemp et al. (2019) may reflect fundamental differences between deliberately established enrichment plantings and naturally assembling seedling communities.

We did find differences by planting configuration for several ecosystem functions, with mixed plantings associated with higher ant abundance. This could be owing to the more variable resources associated with this treatment. For example, tree species differ in structural and functional traits and in the resources they contribute (Butz et al., 2023; Wildermuth et al., 2023). Therefore, diverse tree communities have more heterogeneous vegetation structure and resources (Dantas et al., 2026). Similar effects have also been found in other studies, with ground-ant abundance and ant-mediated seed dispersal increasing alongside tree richness in a tropical biodiversity experiment (Dantas et al., 2024). In the broader literature, experimentally increasing tree richness has been associated with higher herbivorous and predatory arthropod abundance, with this higher diversity contributing to elevated forest productivity (Li et al., 2023). However, these responses are not uniform across arthropod groups (Dantas et al., 2026). For example, in other enrichment planting studies, arthropod groups responded differently to planting structure and type, including Hymenoptera increasing with tree heights, and Hemiptera with heterospecific plantings (Dantas et al., 2026). This likely reflects differences in resource needs and habitat requirements across taxa and functional groups (Cole et al., 2016; Potapov et al., 2020). Our results add to the literature suggesting that mixed plantings may boost biodiversity by increasing habitat and resource diversity.

Soil aggregate stability was also greater in mixed than in conspecific plantings. Gould et al. (2016) similarly found that greater plant diversity increased soil aggregate stability in controlled and long-term field experiments, concluding that this was linked to root traits. Although root traits were not measured in our study, greater diversity in rooting strategies and associated below-ground processes has been a proven mechanism associated with higher aboveground plant diversity in other systems (Ford et al., 2016; Gould et al., 2016) and is therefore likely to be the explanatory factor in our study.

We raise a few potential causes for the contrast between our seedling and ecosystem function results and those from earlier studies on planting configuration. First, survey coverage of the RERTA riparian margins changed over the years, with some years having subsets of the margins. To reduce confounding unsurveyed planting configuration observations with true mortality events in our analysis, we restricted our modelling to the first 5-6 years. Although this reduced the potential risk of misclassification, it also shortened the monitoring period, which may have captured true mortality events related to planting configuration. Time has consistently been a significant factor in mortality for both planting configuration and position in our project, and earlier long-term studies also show neighbourhood effects can operate across multiple years, with their strength varying over time (Hazelwood et al., 2021; Zhu et al., 2015). Therefore, our truncated monitoring period provides evidence for planting-configuration impacts during the first 6 years. Second, our ecosystem surveys focused on three of the six planted native tree species (*Falcataria moluccana*, *Alstonia scholaris*, and *Peronema canescens)*. Although we made this decision to ensure adequate field replicates, we acknowledge we may have masked the ecosystem analysis within the margin of trees that were more impacted during the initial restoration phase. Species are known to vary significantly to their local environment (Comita et al., 2014), and this action may have inadvertently limited our ecosystem surveys to trees that were more robust to the CNDD phenomenon in the field and may underrepresent species that are more sensitive to their neighbours during the initial restoration phase. Therefore, we interpreted our results as a subset of the original planting community.

### Placement within the riparian margin influences seedling performance

Individual seedling mortality varied with position in the riparian margin, with higher mortality closer to the oil palm edge than to the river. This effect likely reflects reduced water availability and climate stability farther from waterways, increasing plant mortality (Allen, 2016; Dala-Corte et al., 2026; Williamson et al., 2020). Other tropical field experiments have also shown that seedling survival and growth differ substantially between riparian margins and upslope, with habitat conditions sometimes explaining more variation in seedling survival than conspecific density (Johnson et al., 2017; O’Brien & Escudero, 2022). Thus, rather than mortality responding uniformly to distance from the river, our results suggest that position within the riparian margin structured seedling survival and that the strength of this spatial pattern depended partly on the surrounding restoration environment.

Planting configuration and distance from the river showed stronger combined than independent effects. Total arthropod abundance declined with increasing distance from the river in conspecific plantings under both restoration treatments, whereas abundance remained comparatively stable across the margin in mixed plantings. Predation showed a similar pattern under Planting Only, declining with distance in conspecific plantings but remaining relatively stable in mixed plantings. Caterpillars showed a different response: under Mature Palms and Planting, abundance increased with distance from the river in mixed plantings but showed no significant spatial trend in conspecific plantings, while neither configuration responded significantly to distance under Planting Only. Thus, planting configuration did not consistently increase arthropod abundance or ecosystem functioning; instead, it altered how ecological responses were expressed across the riparian gradient, with the direction and strength of these effects depending on the wider restoration treatment.

Greater heterogeneity in mixed plantings could provide a broader range of microhabitats and resources across this gradient, potentially buffering associated arthropod communities and predation against changes in conditions as distance from the river increased. Riparian environmental gradients can similarly structure arthropod communities and their functions. Riparian vegetation modifies temperature, humidity and resource availability, which can shape community structure and resources (Allen, 2016). A recent global synthesis found substantial differences between different taxonomic groups, with estimated requirements being narrower for invertebrates than for birds and mammals (Dala-Corte et al., 2026). These studies may help explain the patterns we observed between planting configuration and distance, as environmental conditions changed across riparian margins, but the impact of this gradient may depend on the local tree neighbourhood.

### Application to restoration within tropical agriculture

Our results point towards several practical considerations for restoration within tropical agricultural landscapes. First, mixed-species planting may be preferable to establishing conspecific-only vegetation, resulting in higher predator densities and improved soil conditions. Furthermore, given evidence from the wider literature that succession and competition between trees at different life stages can mask CNDD effects (Comita et al., 2014; Zhu et al., 2015), establishing a diverse first-generation forest may also improve the natural succession of seedlings. However, we acknowledge that this approach may be more logistically challenging and costly for restoration initiatives than replanting with a narrower species range. Furthermore, our results indicate that planting configuration should be considered alongside broader environmental conditions, as conspecific plantings were more disadvantaged under the Planting Only treatment, likely due to higher heat stress and less canopy shade in the Planting Only margin. Therefore, at field sites with harsher climatic conditions and without the option of shade crops, mixed plantings in active restoration may again increase restoration success. Rather than prescribing a single planting configuration across an entire restoration site, this indicates that more diverse planting may be beneficial in areas with more exposed conditions, and that environmental tolerances of species should be considered when determining where to plant them.

These considerations may be particularly important where restoration occurs within landscapes with a long history of intensive agriculture. Active restoration is often recommended when natural regeneration is unlikely because of site conditions, competition from existing vegetation, or limited seed sources (Rodrigues et al., 2011). In our system, riparian margins occur within an oil palm landscape and are isolated from natural forests, limiting the likelihood of natural recolonisation (Luke et al., 2020). Enrichment planting can overcome this dispersal limitation, but our findings suggest that how planted individuals are arranged can be important alongside increasing tree numbers. Evidence from the large-scale Sabah Biodiversity Experiment, in Malaysia, similarly shows that diverse enrichment planting can accelerate aspects of tropical forest structural recovery, supporting species-rich planting strategies in degraded forests (Veryard et al., 2023).

Responses varied among ecosystem metrics, restoration treatments and positions within the margin. So, we caution against assuming our results would be the same in other regions and agricultural systems without further replication or studies with comparable sites and results. Furthermore, because time drove seedling survival, we suggest restoration projects benefit from monitoring planting beyond initial establishment and tracking ecosystem responses as they mature.

Finally, restoration design should account for placement within the restored margin, with plantings near rivers having higher survival than those farther away. This highlights the value of targeting restoration efforts on river margins for establishment success, as well as the wide range of environmental benefits highlighted in other studies (Luke et al., 2019). In addition, several ecosystem responses were shaped by the interaction between planting configuration and distance from the river rather than by either factor alone. This indicates that a planting design may not perform equally well across the full width of a restored margin. Restoration treatments should therefore explicitly incorporate environmental gradients, matching species and neighbourhood composition to variation in moisture, microclimate, and other conditions across riparian margins, rather than applying uniform planting arrangements throughout.

The interconnectedness of several measured responses to restoration treatment further indicates that planting configuration and position cannot be considered independently of the wider restoration environment. Retaining mature palms alongside enrichment planting created different structural and environmental conditions than planting in open margins, likely contributing to higher mortality in the open margin alone and to the distance from the river, which likely increased mortality disproportionately at the oil palm edge. From a restoration perspective, these context-dependent responses indicate that enrichment-planting design should account for both existing vegetation and environmental variation, rather than assuming a single planting configuration will produce consistent outcomes across a site. Combining diverse enrichment plantings with retention of existing vegetation, where appropriate, may increase habitat heterogeneity and provide a broader range of conditions for ecological recovery.

Overall, our findings support diversified rather than uniform enrichment planting; however, we caution against treating mixed planting as superior. If anything, our results show that the effects of planting configuration are context-dependent and can be expressed at different levels, with mixed and conspecific plantings benefiting different ecosystem properties and, for some, being significant only when combined with the wider treatment rather than producing a consistent response.

## References

1. Allen, D. C. (2016). Microclimate modification by riparian vegetation affects the structure and resource limitation of arthropod communities. Ecosphere, 7(2). 10.1002/ecs2.1200

2. Andrews, S. S., Karlen, D. L., & Mitchell, J. P. (2002). A comparison of soil quality indexing methods for vegetable production systems in Northern California. *Agriculture*, Ecosystems and Environment, 90(1). 10.1016/S0167-8809(01)00174-8

3. Anggraini, E., Setiawati, T., Herlinda, S., Irsan, C., Mulawarman, Gofar, N., Muslim, A., & Lau, W. H. (2025). Identification of the nettle caterpillar in smallholding oil palm plantation cultivated on peatland in Ogan Ilir, South Sumatra, Indonesia. Biodiversitas, 26(1). 10.13057/biodiv/d260104

4. Bagchi, R., Swinfield, T., Gallery, R. E., Lewis, O. T., Gripenberg, S., Narayan, L., & Freckleton, R. P. (2010). Testing the Janzen-Connell mechanism: Pathogens cause overcompensating density dependence in a tropical tree. Ecology Letters, 13(10). 10.1111/j.1461-0248.2010.01520.x

5. Barclay, H., Gray, C. L., Luke, S. H., Nainar, A., Snaddon, J. L., & Turner, E. C. (2017). RSPO Manual on Best Management Practices (BMPs) for the Management and Rehabilitation of Riparian Reserves Document Title: RSPO Manual on Best Management Practices (BMPs) for the Management and Rehabilitation of Riparian Reserves. 10.13140/RG.2.2.17011.22561

6. Bouyoucos, G. J. (1962). Hydrometer Method Improved for Making Particle Size Analyses of Soils 1 . Agronomy Journal, 54(5). 10.2134/agronj1962.00021962005400050028x

7. Bray, R. H., & Kurtz, L. T. (1945). Determination of total, organic, and available forms of phosphorus in soils. Soil Science, 59(1). 10.1097/00010694-194501000-00006

8. Bremner, J. M. (1960). Determination of nitrogen in soil by the Kjeldahl method. The Journal of Agricultural Science, 55(1). 10.1017/S0021859600021572

9. Brooks, M. E., Kristensen, K., Benthem, K. J., van Magnusson, A., Berg, C. W., Nielsen, A., Skaug, H. J., Mächler, M., & Bolker, B. M. (2017). glmmTMB Balances Speed and Flexibility Among Packages for Zero-inflated Generalized Linear Mixed Modeling. The R Journal, 9(2), 378. 10.32614/RJ-2017-066

10. Butz, E. M., Schmitt, L. M., Parker, J. D., & Burghardt, K. T. (2023). Positive tree diversity effects on arboreal spider abundance are tied to canopy cover in a forest experiment. Ecology, 104(8). 10.1002/ecy.4116

11. Cole, R. J., Holl, K. D., Zahawi, R. A., Wickey, P., & Townsend, A. R. (2016). Leaf litter arthropod responses to tropical forest restoration. Ecology and Evolution, 6(15). 10.1002/ece3.2220

12. Comita, L. S., Queenborough, S. A., Murphy, S. J., Eck, J. L., Xu, K., Krishnadas, M., Beckman, N., & Zhu, Y. (2014). Testing predictions of the Janzen-Connell hypothesis: A meta-analysis of experimental evidence for distance- and density-dependent seed and seedling survival. Journal of Ecology, 102(4). 10.1111/1365-2745.12232

13. Connell, J. (1971). On the role of natural enemies in preventing competitive exclusion in some marine animals and in rain forest trees. In Dynamics of populations (Vol. 298).

14. Dala-Corte, R. B., Giam, X., & Wilcove, D. S. (2026). Assessing the effectiveness of riparian buffers in protecting biodiversity: a meta-analysis. Nature Communications, 17(1). 10.1038/s41467-026-70191-y

15. Dantas, A., Dantas, T. A. V., Ganade, G., & Fonseca, C. R. (2024). Tree richness affects ground-ant diversity and seed dispersal in a tropical biodiversity– ecosystem function experiment. Oikos, 2024(10). 10.1111/oik.10623

16. Dantas, T. A. V., Dantas, A., Ganade, G., & Fonseca, C. R. (2026). Ground-dwelling arthropod community structure responds to an experimental gradient of tropical tree diversity. Oecologia, 208(1). 10.1007/s00442-025-05838-6

17. dos Santos, W. L., de Oliveira, C. D. C., & Durigan, G. (2025). Enrichment planting under restored riparian forests: neither for all species nor for all situations. Biological Conservation, 307, 111205. 10.1016/J.BIOCON.2025.111205

18. Eijkelkamp, R. (2022). Wet sieving apparatus for disturbed samples.

19. FAO. (2023a). FAOSTAT on-line Database. Food and Agriculture Organization of the United Nations.

20. FAO. (2023b). Standard operating procedure for soil moisture content by gravimetric method. http://www.wipo.int/amc/en/mediation/rules

21. Fichtner, A., Härdtle, W., Bruelheide, H., Kunz, M., Li, Y., & Von Oheimb, G. (2018). Neighbourhood interactions drive overyielding in mixed-species tree communities. Nature Communications, 9(1). 10.1038/s41467-018-03529-w

22. Ford, H., Garbutt, A., Ladd, C., Malarkey, J., & Skov, M. W. (2016). Soil stabilization linked to plant diversity and environmental context in coastal wetlands. Journal of Vegetation Science, 27(2). 10.1111/jvs.12367

23. Gibbs, H. K., Ruesch, A. S., Achard, F., Clayton, M. K., Holmgren, P., Ramankutty, N., & Foley, J. A. (2010). Tropical forests were the primary sources of new agricultural land in the 1980s and 1990s. Proceedings of the National Academy of Sciences of the United States of America, 107(38). 10.1073/pnas.0910275107

24. Gould, I. J., Quinton, J. N., Weigelt, A., De Deyn, G. B., & Bardgett, R. D. (2016). Plant diversity and root traits benefit physical properties key to soil function in grasslands. In Ecology letters (Vol. 19, Number 9). 10.1111/ele.12652

25. Hazelwood, K., Beck, H., & Paine, C. E. T. (2021). Negative density dependence in the mortality and growth of tropical tree seedlings is strong, and primarily caused by fungal pathogens. Journal of Ecology, 109(4). 10.1111/1365-2745.13615

26. Janzen, D. H. (1970). Herbivores and the Number of Tree Species in Tropical Forests. The American Naturalist, 104(940). 10.1086/282687

27. Jevon, F. V., De La Cruz, D., LaManna, J. A., Lang, A. K., Orwig, D. A., Record, S., Kouba, P. V., Ayres, M. P., & Matthes, J. H. (2022). Experimental and observational evidence of negative conspecific density dependence in temperate ectomycorrhizal trees. Ecology, 103(11). 10.1002/ecy.3808

28. Johnson, D. J., Condit, R., Hubbell, S. P., & Comita, L. S. (2017). Abiotic niche partitioning and negative density dependence drive tree seedling survival in a tropical forest. Proceedings of the Royal Society B: Biological Sciences, 284(1869). 10.1098/rspb.2017.2210

29. Kaiser, H. F. (1960). The Application of Electronic Computers to Factor Analysis. Educational and Psychological Measurement, 20(1). 10.1177/001316446002000116

30. Li, Y., Schmid, B., Schuldt, A., Li, S., Wang, M. Q., Fornoff, F., Staab, M., Guo, P. F., Anttonen, P., Chesters, D., Bruelheide, H., Zhu, C. D., Ma, K., & Liu, X. (2023). Multitrophic arthropod diversity mediates tree diversity effects on primary productivity. Nature Ecology and Evolution, 7(6). 10.1038/s41559-023-02049-1

31. Luke, S. H., Advento, A. D., Aryawan, A. A. K., Adhy, D. N., Ashton-Butt, A., Barclay, H., Dewi, J. P., Drewer, J., Dumbrell, A. J., Edi Eycott, A. E., Harianja, M. F., Hinsch, J. K., Hood, A. S. C., Kurniawan, C., Kurz, D. J., Mann, D. J., Matthews Nicholass, K. J., Naim, M., … Turner, E. C. (2020). Managing Oil Palm Plantations More Sustainably: Large-Scale Experiments Within the Biodiversity and Ecosystem Function in Tropical Agriculture (BEFTA) Programme. Frontiers in Forests and Global Change, 2. 10.3389/ffgc.2019.00075

32. Luke, S. H., Slade, E. M., Gray, C. L., Annammala, K. V., Drewer, J., Williamson, J., Agama, A. L., Ationg, M., Mitchell, S. L., Vairappan, C. S., & Struebig, M. J. (2019). Riparian buffers in tropical agriculture: Scientific support, effectiveness and directions for policy. Journal of Applied Ecology, 56(1), 85–92. 10.1111/1365-2664.13280

33. Meli, P., Lozano-Báez, S. E., Rother, D. C., Bustamante-Sánchez, M. A., Zeller, M., & Rodrigues, R. R. (2026). Enrichment planting in tropical and subtropical forest restoration: patterns, practices, and knowledge gaps. In Restoration Ecology. John Wiley and Sons Inc. 10.1111/rec.70485

34. Motsara, M. R. ., & Roy, R. N. . (2008). Guide to laboratory establishment for plant nutrient analysis (Vol. 19). Food and Agriculture Organization of the United Nations.

35. O’Brien, M. J., & Escudero, A. (2022). Topography in tropical forests enhances growth and survival differences within and among species via water availability and biotic interactions. Functional Ecology, 36(3). 10.1111/1365-2435.13977

36. Pendrill, F., Gardner, T. A., Meyfroidt, P., Persson, U. M., Adams, J., Azevedo, T., Lima, M. G. B., Baumann, M., Curtis, P. G., De Sy, V., Garrett, R., Godar, J., Goldman, E. D., Hansen, M. C., Heilmayr, R., Herold, M., Kuemmerle, T., Lathuillière, M. J., Ribeiro, V., … West, C. (2022). Disentangling the numbers behind agriculture-driven tropical deforestation. In Science (Vol. 377, Number 6611). 10.1126/science.abm9267

37. Posit team. (2023). RStudio: Integrated Development Environment for R. http://www.posit.co/

38. Potapov, A. M., Dupérré, N., Jochum, M., Dreczko, K., Klarner, B., Barnes, A. D., Krashevska, V., Rembold, K., Kreft, H., Brose, U., Widyastuti, R., Harms, D., & Scheu, S. (2020). Functional losses in ground spider communities due to habitat structure degradation under tropical land-use change. Ecology, 101(3). 10.1002/ecy.2957

39. Ramage, B. S., Johnson, D. J., Gonzalez-Akre, E., McShea, W. J., Anderson-Teixeira, K. J., Bourg, N. A., & Clay, K. (2017). Sapling growth rates reveal conspecific negative density dependence in a temperate forest. Ecology and Evolution, 7(19). 10.1002/ece3.3298

40. Rodrigues, R. R., Gandolfi, S., Nave, A. G., Aronson, J., Barreto, T. E., Vidal, C. Y., & Brancalion, P. H. S. (2011). Large-scale ecological restoration of high-diversity tropical forests in SE Brazil. Forest Ecology and Management, 261(10), 1605– 1613. 10.1016/j.foreco.2010.07.005

41. Schneider, C. A., Rasband, W. S., & Eliceiri, K. W. (2012). NIH Image to ImageJ: 25 years of image analysis. In Nature Methods (Vol. 9, Number 7). 10.1038/nmeth.2089

42. Schweizer, D., Gilbert, G. S., & Holl, K. D. (2013). Phylogenetic ecology applied to enrichment planting of tropical native tree species. Forest Ecology and Management, 297. 10.1016/j.foreco.2013.02.012

43. Shokr, M. S., Jalhoum, M. E. M., Sayed, A. S. A., Saeed, M., Rebouh, N. Y., Mohamed, E. S., Yousif, I. A. H., & Abdelhameed, H. H. (2025). Assessment of soil quality in arid zones using principal component analysis and GIS-based modeling. PLOS ONE, 20(12 December). 10.1371/journal.pone.0337063

44. Teuscher, M., Gérard, A., Brose, U., Buchori, D., Clough, Y., Ehbrecht, M., Hölscher, D., Irawan, B., Sundawati, L., Wollni, M., & Kreft, H. (2016). Experimental biodiversity enrichment in oil-palm-dominated landscapes in Indonesia. Frontiers in Plant Science, 7(OCTOBER 2016). 10.3389/fpls.2016.01538

45. Veryard, R., Wu, J., O’Brien, M. J., Anthony, R., Both, S., Burslem, D. F. R. P., Chen, B., Cagigal, E. F. M., Godfray, C. H. J., Godoong, E., Liang, S., Saner, P., Schmid, B., Wai, Y. S., Xie, J., Reynolds, G., & Hector, A. (2023). Positive effects of tree diversity on tropical forest restoration in a field-scale experiment. Science Advances, 9(37). 10.1126/sciadv.adf0938

46. Walkley, A., & Black, I. A. (1934). An examination of the degtjareff method for determining soil organic matter, and a proposed modification of the chromic acid titration method. Soil Science, 37(1). 10.1097/00010694-193401000-00003

47. Webb, C. O., & Peart, D. R. (1999). Seedling density dependence promotes coexistence of Bornean rain forest trees. Ecology, 80(6). 10.1890/0012-9658(1999)080[2006:SDDPCO]2.0.CO;2

48. Wildermuth, B., Dönges, C., Matevski, D., Penanhoat, A., Seifert, C. L., Seidel, D., Scheu, S., & Schuldt, A. (2023). Tree species identity, canopy structure and prey availability differentially affect canopy spider diversity and trophic composition. Oecologia, 203(1–2). 10.1007/s00442-023-05447-1

49. Williamson, J., Slade, E. M., Luke, S. H., Swinfield, T., Chung, A. Y. C., Coomes, D. A., Heroin, H., Jucker, T., Lewis, O. T., Vairappan, C. S., Rossiter, S. J., & Struebig, M. J. (2020). Riparian buffers act as microclimatic refugia in oil palm landscapes. Journal of Applied Ecology, 58(2), 431–442. 10.1111/1365-2664.13784

50. Xu, Y., Yu, L., Ciais, P., Li, W., Santoro, M., Yang, H., & Gong, P. (2022). Recent expansion of oil palm plantations into carbon-rich forests. Nature Sustainability, 5(7). 10.1038/s41893-022-00872-1

51. Yamada, T., Watanabe, K., Okuda, T., Sugimoto, T., & Noor Azlin, Y. (2016). Growth and survival of trees planted in an oil palm plantation: Implications for restoration of biodiversity. Journal of Tropical Forest Science, 28(1).

52. Zemp, D. C., Ehbrecht, M., Seidel, D., Ammer, C., Craven, D., Erkelenz, J., Irawan, B., Sundawati, L., Hölscher, D., & Kreft, H. (2019). Mixed-species tree plantings enhance structural complexity in oil palm plantations. Agriculture, Ecosystems and Environment, 283. 10.1016/j.agee.2019.06.003

53. Zemp, D. C., Gérard, A., Hölscher, D., Ammer, C., Irawan, B., Sundawati, L., Teuscher, M., & Kreft, H. (2019). Tree performance in a biodiversity enrichment experiment in an oil palm landscape. Journal of Applied Ecology, 56(10), 2340– 2352. 10.1111/1365-2664.13460

54. Zemp, D. C., Guerrero-Ramirez, N., Brambach, F., Darras, K., Grass, I., Potapov, A., Röll, A., Arimond, I., Ballauff, J., Behling, H., Berkelmann, D., Biagioni, S., Buchori, D., Craven, D., Daniel, R., Gailing, O., Ellsäßer, F., Fardiansah, R., Hennings, N., … Kreft, H. (2023). Tree islands enhance biodiversity and functioning in oil palm landscapes. Nature, 618(7964). 10.1038/s41586-023-06086-5

55. Zhu, Y., Comita, L. S., Hubbell, S. P., & Ma, K. (2015). Conspecific and phylogenetic density-dependent survival differs across life stages in a tropical forest. Journal of Ecology, 103(4). 10.1111/1365-2745.12414

