## Supplementary Figure S1 for "Planting set-up and local environmental conditions affect outcomes in tropical agricultural restoration"

### Supplementary Material

**Supplementary Table S1 | Mealworm predation rate**

A) **Overall predation rate:** General Linear Mixed Model with fixed effects (group) and their interactions. Treatment includes two active restorations (Mature Palms and Planting and Planting Only), two Planting types for the enrichment plantings (conspecific seedlings all the same species, mixed seedlings), at various distances within the margin (meters from the river edge), and the interactions between all the different fixed effects.

| Group | *Chisq* | *df* | *p* |
| --- | --- | --- | --- |
| Treatment | 20.9497 | 1 | 4.715e-06 |
| Planting Type | 9.5487 | 1 | 0.002001 |
| Distance | 0.3422 | 1 | 0.558574 |
| Treatment: Planting | 3.0380 | 1 | 0.081334 |
| Treatment: Distance | 1.8216 | 1 | 0.177119 |
| Planting: Distance | 5.3386 | 1 | 0.020858 |
| Treatment: Planting: Distance | 1.7458 | 1 | 0.186411 |

B) **Pairwise comparisons:** estimated marginal means (EMMs) and estimated marginal trends for mealworm predation rate from Table S2 (A). Treatment and planting contrasts are presented as odds ratios (OR). Comparisons involving distance represent differences in estimated slopes (β) because distance was fitted as a continuous variable. OR = odds ratio; β = difference between slopes; SE = standard error. Distance was included as a continuous predictor; therefore, comparisons involving distance were obtained using emtrends.

| Effect | Conditioning variable | Contrast | Estimate | SE | z | *p* |
| --- | --- | --- | --- | --- | --- | --- |
| Treatment | — | MP + P / P Only | OR = 3.47 | 0.986 | 4.381 | <0.0001 |
| Planting Type | — | Conspecific / Mixed | OR = 1.60 | 0.530 | 1.416 | 0.157 |
| Distance slope by planting | MP + P | Conspecific − Mixed | β = −0.0230 | 0.0339 | −0.680 | 0.497 |
| Distance slope by planting | P Only | Conspecific − Mixed | β = −0.0844 | 0.0326 | −2.589 | 0.0096 |

**Supplementary Table S2. Arthropod abundance counts**

(A) **Overall model effects:** General Linear Mixed Model with fixed effects (Model Term) and their interactions. Treatment includes two active restorations (Mature Palms and Planting and Planting Only), two Planting types for the enrichment plantings (conspecific seedlings all the same species, mixed seedlings), at various distances within the margin (meters from the river edge), and the interactions between all the different fixed effects on the overall arthropod abundance and individual predator (Weaver Ants, Total Ants, Spiders) and pest groups (Overall caterpillar, nettle caterpillars, and bagworm caterpillars). † Bagworm and Nettle caterpillar abundance was modelled additively (Treatment + Planting type + Distance) rather than using the full factorial interaction structure because observations were too sparse to support estimation of the interaction terms. All other models included the full Treatment × Planting type × Distance interaction. Bold indicates *p* < 0.05.

| Arthropod group | Model term | χ² | *df* | *p* |
| --- | --- | --- | --- | --- |
| Total arthropods | Treatment | 0.677 | 1 | 0.411 |
|  | Planting type | 2.071 | 1 | 0.150 |
|  | Distance | 3.235 | 1 | 0.072 |
|  | Treatment × Planting type | 0.177 | 1 | 0.674 |
|  | Treatment × Distance | 1.020 | 1 | 0.312 |
|  | Planting type × Distance | 12.405 | 1 | <0.001 |
|  | Treatment × Planting type × Distance | 0.005 | 1 | 0.947 |
| Weaver ants | Treatment | 0.085 | 1 | 0.771 |
|  | Planting type | 0.557 | 1 | 0.456 |
|  | Distance | 0.387 | 1 | 0.534 |
|  | Treatment × Planting type | 3.771 | 1 | 0.052 |
|  | Treatment × Distance | 1.564 | 1 | 0.211 |
|  | Planting type × Distance | 0.063 | 1 | 0.803 |
|  | Treatment × Planting type × Distance | 2.984 | 1 | 0.084 |
| Total ants | Treatment | 2.054 | 1 | 0.152 |
|  | Planting type | 12.288 | 1 | <0.001 |
|  | Distance | 3.907 | 1 | 0.048 |
|  | Treatment × Planting type | 2.074 | 1 | 0.150 |
|  | Treatment × Distance | 0.710 | 1 | 0.399 |
|  | Planting type × Distance | 2.532 | 1 | 0.112 |
|  | Treatment × Planting type × Distance | 3.734 | 1 | 0.053 |
| Spiders | Treatment | 1.633 | 1 | 0.201 |
|  | Planting type | 3.721 | 1 | 0.054 |
|  | Distance | 0.488 | 1 | 0.485 |
|  | Treatment × Planting type | 0.351 | 1 | 0.553 |
|  | Treatment × Distance | 0.601 | 1 | 0.438 |
|  | Planting type × Distance | 0.478 | 1 | 0.489 |
|  | Treatment × Planting type × Distance | 0.108 | 1 | 0.743 |
| Caterpillars | Treatment | 11.659 | 1 | <0.001 |
|  | Planting type | 0.858 | 1 | 0.354 |
|  | Distance | 1.162 | 1 | 0.281 |
|  | Treatment × Planting type | 2.905 | 1 | 0.088 |
|  | Treatment × Distance | 0.190 | 1 | 0.663 |
|  | Planting type × Distance | 1.771 | 1 | 0.183 |
|  | Treatment × Planting type × Distance | 5.678 | 1 | 0.017 |
| Bagworm caterpillars† | Treatment | 2.7133 | 1 | 0.1 |
|  | Planting type | 0.006 | 1 | 0.937 |
|  | Distance | 3.743 | 1 | 0.053 |
| Nettle caterpillars† | Treatment | 0.000 | 1 | 1.000 |
|  | Planting type | 0.031 | 1 | 0.860 |
|  | Distance | 1.377 | 1 | 0.241 |

**(B) Post-hoc comparisons:** estimated marginal means and marginal trends associated with significant effects in the overall models (Supplementary Table S3 A). Response ratios are reported for count-model contrasts, while β represents estimated distance slopes or differences between distance slopes. Wald χ² statistics. We calculated estimated marginal means and trends using emmeans. Distance was fitted as a continuous predictor; consequently, effects involving distance were evaluated using emtrends() rather than treating distance as a categorical variable. Response ratios <1 indicate lower predicted abundance in the numerator group relative to the denominator group. Bold indicates *p* < 0.05.

| B1. Total arthropod abundance: estimated distance trends | | | | | |
| --- | --- | --- | --- | --- | --- |
| Treatment | Planting type | Distance slope (β) | SE | 95% CI | - |
| MP + P | Conspecific | −0.0831 | 0.0393 | −0.1602 to −0.0061 | - |
| P Only | Conspecific | −0.1161 | 0.0292 | −0.1733 to −0.0590 | - |
| MP + P | Mixed | 0.0246 | 0.0193 | −0.0132 to 0.0624 | - |
| P Only | Mixed | −0.0041 | 0.0350 | −0.0727 to 0.0645 | - |
| B2. Total arthropod abundance: pairwise comparisons of distance slopes | | | | | |
| Treatment | Contrast | Difference in slopes (β) | SE | z | *p* |
| MP + P | Conspecific − Mixed | −0.108 | 0.0428 | −2.516 | 0.012 |
| P Only | Conspecific − Mixed | −0.112 | 0.0471 | −2.380 | 0.017 |
| B3. Total ant abundance: planting-type comparison | | | | | |
| Contrast | Ratio | SE | z | *p* | - |
| Conspecific / Mixed | 0.470 | 0.109 | −3.260 | 0.0011 | - |
| B4. Caterpillar abundance: treatment comparison | | | | | |
| Contrast | Ratio | SE | z | *p* | - |
| MP + P / P Only | 0.381 | 0.102 | −3.598 | <0.001 | - |

Supplementary Table S3 | Leaf litter trap contents: herbivory

A) **Overall model effects:** General Linear Mixed Model for effects of two active restoration treatments (Mature Palms and Planting and Planting Only), planting type (Conspecific planting with the same species, mixed seedling species planted together), distance (meters from the river), and their interactions. Bold indicates *p* < 0.05.

| Model term | Chisq | Df | P value |
| --- | --- | --- | --- |
| Treatment | 16.677 | 1 | 4.431e-05 |
| Planting type | 1.196 | 1 | 0.27415 |
| Distance | 2.219 | 1 | 0.13635 |
| Treatment: Planting | 0.684 | 1 | 0.40818 |
| Treatment: Distance | 3.05 | 1 | 0.08073 |
| Planting: Distance | 0.534 | 1 | 0.46489 |
| Treatment: Planting: Distance | 1.828 | 1 | 0.17639 |

B) **Post hoc comparison for treatment level**

| Treatment | emmeans | SE | Lower CI | Upper CI |
| --- | --- | --- | --- | --- |
| MP + P | -0.759 | 0.108 | -0.969 | -0.544 |
| P Only | -0.19 | 0.0981 | -0.383 | 0.00209 |
| Contrast | estimate | SE | T ratio | P value |
| MP + P – P only | -0.566 | 0.135 | -4.195 | <.0001 |

**Supplementary S4 | Leaf Litter Trap Contents: Biomass**

A) **Overall Model effects:** General Linear Mixed Model for effects of two active restoration treatments (Mature Palms and Planting and Planting Only), planting type (Conspecific planting with the same species, mixed seedling species planted together), distance (meters from the river), and their interactions. Bold indicates *p* < 0.05.

| Model term | Chisq | Df | P value |
| --- | --- | --- | --- |
| Treatment | 73.0596 | 1 | <2e-16 *** |
| Planting type | 1.236 | 1 | 0.2662 |
| Distance | 0.6221 | 1 | 0.4303 |
| Category | 231.2645 | 4 | <2e-16 *** |
| Treatment: Planting | 1.0993 | 1 | 0.2944 |
| Treatment: Distance | 1.9077 | 1 | 0.1672 |
| Planting: Distance | 2.074 | 1 | 0.1498 |
| Treatment: Planting: Distance | 0.2585 | 1 | 0.6112 |

B) **Post hoc comparison:** on significant terms from Supplementary S6 (A), overall ANOVA of the leaf litter trap contents analysed by biomass.

| B1. Post hoc comparison: Treatment | | | | |
| --- | --- | --- | --- | --- |
| Treatment | response | SE | Lower CI | Upper CI |
| MP + P | 0.0783 | 0.0168 | 0.0514 | 0.119 |
| P Only | 0.2064 | 0.0455 | 0.1341 | 0.318 |
| Contrast | ratio | SE | Z ratio | P value |
| MP + P – P only | 0.379 | 0.0426 | -8.638 | <.0001 |
| B2. Post Hoc comparison: Category | | | | |
| Category | ratio | SE | Z ratio | P value |
| Leaves | 0.379 | 0.0426 | -8.638 | <.0001 |
| Wood | 0.379 | 0.0426 | -8.638 | <.0001 |
| Seeds | 0.379 | 0.0426 | -8.638 | <.0001 |
| Flowers | 0.379 | 0.0426 | -8.638 | <.0001 |
| Fruit | 0.379 | 0.0426 | -8.638 | <.0001 |

**Supplementary Table S5 | Soil Chemical Properties**

A) **Overall model effects:** General Linear Mixed Model with fixed effects (Model Term) and their interactions. Treatment includes two active restorations (Mature Palms and Planting and Planting Only), two Planting types for the enrichment plantings (conspecific seedlings all the same species, mixed seedlings), at various distances within the margin (meters from the river edge), and the interactions between all the different fixed effects on the soil chemical properties. Bold indicates *p* < 0.05.

| Soil property | Model term | Test statistic | df | *p* |
| --- | --- | --- | --- | --- |
| CEC | Treatment | 0.007 | 1 | 0.933 |
|  | Planting type | 0.769 | 1 | 0.381 |
|  | Distance | 0.054 | 1 | 0.816 |
|  | Treatment × Planting type | 0.266 | 1 | 0.606 |
|  | Treatment × Distance | 0.009 | 1 | 0.925 |
|  | Planting type × Distance | 1.672 | 1 | 0.196 |
|  | Treatment × Planting type × Distance | 0.546 | 1 | 0.460 |
| Organic carbon | Treatment | 10.899 | 1 | <0.001 |
|  | Planting type | 7.925 | 1 | 0.005 |
|  | Distance | 16.564 | 1 | <0.001 |
|  | Treatment × Planting type | 12.351 | 1 | <0.001 |
|  | Treatment × Distance | 1.388 | 1 | 0.239 |
|  | Planting type × Distance | 7.421 | 1 | 0.006 |
|  | Treatment × Planting type × Distance | 0.940 | 1 | 0.332 |
| Phosphorus | Treatment | 0.837 | 1 | 0.360 |
|  | Planting type | 0.910 | 1 | 0.340 |
|  | Distance | 1.022 | 1 | 0.312 |
|  | Treatment × Planting type | 0.001 | 1 | 0.972 |
|  | Treatment × Distance | 1.737 | 1 | 0.188 |
|  | Planting type × Distance | 1.581 | 1 | 0.209 |
|  | Treatment × Planting type × Distance | 0.375 | 1 | 0.540 |
| Total nitrogen | Treatment | 1.222 | 1 | 0.269 |
|  | Planting type | 0.552 | 1 | 0.458 |
|  | Distance | 0.262 | 1 | 0.609 |
|  | Treatment × Planting type | 0.050 | 1 | 0.824 |
|  | Treatment × Distance | 1.240 | 1 | 0.266 |
|  | Planting type × Distance | 1.549 | 1 | 0.213 |
|  | Treatment × Planting type × Distance | <0.001 | 1 | 0.992 |
| pH | Treatment | 0.004 | 1 | 0.948 |
|  | Planting type | 0.126 | 1 | 0.722 |
|  | Distance | 0.015 | 1 | 0.904 |
|  | Treatment × Planting type | 1.918 | 1 | 0.166 |
|  | Treatment × Distance | 0.016 | 1 | 0.900 |
|  | Planting type × Distance | 0.020 | 1 | 0.887 |
|  | Treatment × Planting type × Distance | 0.695 | 1 | 0.404 |

(B) **Post-hoc comparisons:** results from the GLMM ANOVA are in Supplementary Table S4 for significant results. Bold indicates *p* < 0.05.

| B1. Organic carbon | | | | | | |
| --- | --- | --- | --- | --- | --- | --- |
| Effect | Conditioning variable | Contrast | Estimate | SE | z | *p* |
| Treatment | Overall | MP + P − P Only | 0.0906 | 0.0289 | 3.133 | 0.0017 |
| Planting type | Overall | Conspecific − Mixed | −0.0521 | 0.0291 | −1.790 | 0.0735 |
| Treatment | Conspecific | MP + P − P Only | −0.0048 | 0.0398 | −0.121 | 0.9036 |
| Treatment | Mixed | MP + P − P Only | 0.1860 | 0.0425 | 4.380 | <0.0001 |
| B2. Organic-carbon distance slopes.  β represents the estimated change in organic carbon for a one-unit increase in distance from the river. | | | | | | |
| Treatment | Planting Type | Distance slope (β) | SE | 95% CI | z | *p* |
| MP + P | Conspecific | −0.0021 | 0.0022 | −0.0065 to 0.0022 | −0.951 | 0.342 |
| P Only | Conspecific | −0.0015 | 0.0032 | −0.0078 to 0.0048 | −0.454 | 0.650 |
| MP + P | Mixed | −0.0111 | 0.0023 | −0.0157 to −0.0066 | −4.804 | <0.0001 |
| P Only | Mixed | −0.0051 | 0.0033 | −0.0116 to 0.0015 | −1.521 | 0.128 |
| B3. Overall organic-carbon distance trend | | | | | | |
| Effect | Distance slope (β) | SE | 95% CI | z | *p* |  |
| Distance | −0.00494 | 0.00139 | −0.00767 to −0.00221 | −3.543 | 0.0004 |  |

**Supplementary Table S6| Soil physical properties**

**(A) Overall model effects:** General Linear Mixed Model for effects of two active restoration treatments (Mature Palms and Planting and Planting Only), planting type (Conspecific planting with the same species, mixed seedling species planted together), distance (meters from the river), and their interactions on soil physical properties. Bold indicates *p* < 0.05.

| Soil property | Model term | χ² | df | *p* |
| --- | --- | --- | --- | --- |
| A-horizon depth | Treatment | 5.481 | 1 | 0.019 |
|  | Planting type | 0.049 | 1 | 0.825 |
|  | Distance | 0.317 | 1 | 0.573 |
|  | Treatment × Planting type | 0.314 | 1 | 0.575 |
|  | Treatment × Distance | 1.074 | 1 | 0.300 |
|  | Planting type × Distance | 0.925 | 1 | 0.336 |
|  | Treatment × Planting type × Distance | 0.754 | 1 | 0.385 |
| Aggregate stability | Treatment | 0.735 | 1 | 0.391 |
|  | Planting type | 4.544 | 1 | 0.033 |
|  | Distance | 0.318 | 1 | 0.573 |
|  | Treatment × Planting type | 1.533 | 1 | 0.216 |
|  | Treatment × Distance | 0.008 | 1 | 0.929 |
|  | Planting type × Distance | 1.224 | 1 | 0.269 |
|  | Treatment × Planting type × Distance | 1.859 | 1 | 0.173 |
| Friable soil (%) | Treatment | 2.136 | 1 | 0.144 |
|  | Planting type | 0.372 | 1 | 0.542 |
|  | Distance | 2.065 | 1 | 0.151 |
|  | Treatment × Planting type | 10.128 | 1 | 0.0015 |
|  | Treatment × Distance | 5.083 | 1 | 0.024 |
|  | Planting type × Distance | 1.294 | 1 | 0.255 |
|  | Treatment × Planting type × Distance | 0.460 | 1 | 0.498 |
| Bulk density | Treatment | 12.059 | 1 | <0.001 |
|  | Planting type | 2.138 | 1 | 0.144 |
|  | Distance | 0.540 | 1 | 0.462 |
|  | Treatment × Planting type | 1.771 | 1 | 0.183 |
|  | Treatment × Distance | 2.408 | 1 | 0.121 |
|  | Planting type × Distance | 0.109 | 1 | 0.741 |
|  | Treatment × Planting type × Distance | 4.000 | 1 | 0.045 |

B) **Post-hoc comparisons, estimated marginal means, and distance trends for soil physical properties following significant effects in the fitted models.** Estimates are presented on the model scale. MP + P = enrichment planting with mature palms retained; P Only = enrichment planting with mature palms removed; SE = standard error; CI = confidence interval; β = estimated slope with increasing distance from the river edge. Significant post-hoc comparisons (*p* < 0.05) are shown in bold.

| Soil property / comparison | Condition | Contrast or estimate | Estimate / β | SE | df | *t* | *p* | 95% CI |
| --- | --- | --- | --- | --- | --- | --- | --- | --- |
| A. A-horizon depth | | | | | | | | |
| Treatment contrast | — | MP + P − P Only | −0.914 | 0.397 | 61 | −2.299 | 0.025 | — |
| Estimated marginal mean | MP + P | Estimated depth | 3.58 | 0.282 | — | — | — | 3.02–4.15 |
| Estimated marginal mean | P Only | Estimated depth | 4.49 | 0.280 | — | — | — | 3.93–5.05 |
| B. Aggregate stability | | | | | | | | |
| Planting-type contrast | — | Conspecific − Mixed | −6.49 | 3.10 | 61 | −2.091 | 0.041 | — |
| Estimated marginal mean | Conspecific | Estimated aggregate stability | 81.9 | 2.18 | — | — | — | 77.5–86.2 |
| Estimated marginal mean | Mixed | Estimated aggregate stability | 88.3 | 2.21 | — | — | — | 83.9–92.8 |
| C. Friable soil — Treatment × Planting type | | | | | | | | |
| Treatment contrast | Conspecific | MP + P − P Only | 3.37 | 2.65 | 62 | 1.272 | 0.208 | — |
| Treatment contrast | Mixed | MP + P − P Only | −8.41 | 2.63 | 62 | −3.199 | 0.0022 | — |
| Estimated marginal mean | MP + P × Conspecific | Estimated friable soil (%) | 11.63 | 1.92 | — | — | — | 7.79–15.50 |
| Estimated marginal mean | P Only × Conspecific | Estimated friable soil (%) | 8.27 | 1.93 | — | — | — | 4.42–12.10 |
| Estimated marginal mean | MP + P × Mixed | Estimated friable soil (%) | 6.85 | 1.91 | — | — | — | 3.04–10.70 |
| Estimated marginal mean | P Only × Mixed | Estimated friable soil (%) | 15.26 | 1.93 | — | — | — | 11.41–19.10 |
| D. Friable soil — Treatment × Distance | | | | | | | | |
| Distance trend | MP + P | Distance slope | −0.288 | 0.117 | — | — | — | −0.522 to −0.055 |
| Distance trend | P Only | Distance slope | 0.153 | 0.149 | — | — | — | −0.145 to 0.452 |
| Slope contrast | — | MP + P − P Only | −0.442 | 0.189 | 62 | −2.341 | 0.0225 | — |
| E. Bulk density — Treatment | | | | | | | | |
| Treatment contrast | — | MP + P − P Only | −0.168 | 0.0498 | 66 | −3.362 | 0.0013 | — |
| F. Bulk density — distance trends by Treatment × Planting type | | | | | | | | |
| Distance trend | MP + P × Conspecific | Distance slope | 0.00350 | 0.00440 | — | — | — | −0.00530 to 0.0123 |
| Distance trend | P Only × Conspecific | Distance slope | 0.00219 | 0.00450 | — | — | — | −0.00680 to 0.0112 |
| Distance trend | MP + P × Mixed | Distance slope | −0.00577 | 0.00409 | — | — | — | −0.01395 to 0.00240 |
| Distance trend | P Only × Mixed | Distance slope | **0.01155** | 0.00551 | — | — | — | **0.00053 to 0.0226** |
| Slope contrast | Conspecific | MP + P − P Only | 0.00131 | 0.00629 | 62 | 0.208 | 0.836 | — |
| Slope contrast | Mixed | **MP + P − P Only** | **−0.01732** | 0.00687 | 62 | −2.523 | **0.014** | — |

#### Supplementary Table S7 | Cumulative variance from Principal

**Component Analysis:**

Eigenvalues and the proportion of variance explained by the first four principal components (PCs) from the principal component analysis (PCA) of soil physicochemical properties. Cumulative variance represents the total proportion of variation explained by successive principal components.

| PC axis | Eigenvalue | Percent variance | Cumulative variance |
| --- | --- | --- | --- |
| comp 1 | 7.335462 | 33.34301 | 33.34301 |
| comp 2 | 5.10398 | 23.19991 | 56.54292 |
| comp 3 | 2.267408 | 10.3064 | 66.84932 |
| comp 4 | 1.441324 | 6.551472 | 73.40079 |

Table S8 | Principal component analysis (PCA) loadings for soil physicochemical variables contributing to the first five principal components (PC).

Only variables with absolute loadings ≥ 0.50 are shown. Absolute loading (Abs Loading), percentage contribution to each principal component (Contribution), and squared cosine values (Cos²), representing the quality of variable representation on each principal component, are presented. Variables include Org_Carbon = organic carbon; HCl25_K = 25% HCl-extractable potassium; CEC = cation exchange capacity; Exch_Ca = exchangeable calcium; Exch_Mg = exchangeable magnesium; Exch_K, exchangeable potassium; Sum_Bases, sum of exchangeable base cations; Base_Saturation = base saturation; pH_H2O = soil pH measured in water (H₂O); pH_KCl = soil pH measured in potassium chloride (KCl); Exch_H = exchangeable hydrogen; Exch_Al = exchangeable aluminium; Carbon.Nitrogen = carbon-to-nitrogen (C:N) ratio; HCl25_P = 25% HCl-extractable phosphorus; P_Bray1 = Bray-1 extractable phosphorus; T_Nitrogen = total nitrogen

| PC axis | Variable | Loading | Abs Loading | Contribution | Cos2 |
| --- | --- | --- | --- | --- | --- |
| Dim.1 | HCl25_P | 0.661764 | 0.661764 | 5.970056 | 0.437931 |
| Dim.1 | HCl25_K | 0.751296 | 0.751296 | 7.694747 | 0.564445 |
| Dim.1 | HCl25_Mg | 0.843436 | 0.843436 | 9.697875 | 0.711384 |
| Dim.1 | P_Bray1 | 0.579378 | 0.579378 | 4.576115 | 0.335679 |
| Dim.1 | Exch_Ca | 0.917163 | 0.917163 | 11.46741 | 0.841188 |
| Dim.1 | Exch_Mg | 0.867467 | 0.867467 | 10.25837 | 0.752499 |
| Dim.1 | Exch_K | 0.87794 | 0.87794 | 10.50757 | 0.770779 |
| Dim.1 | Exch_Na | 0.612513 | 0.612513 | 5.114498 | 0.375172 |
| Dim.1 | Sum_Bases | 0.952783 | 0.952783 | 12.37543 | 0.907795 |
| Dim.1 | Base_Saturation | 0.714983 | 0.714983 | 6.968893 | 0.511201 |
| Dim.2 | Water.Content | 0.626867 | 0.626867 | 7.699138 | 0.392962 |
| Dim.2 | Texture_Sand | -0.53525 | 0.535246 | 5.613036 | 0.286488 |
| Dim.2 | pH_H2O | -0.7955 | 0.795495 | 12.39842 | 0.632813 |
| Dim.2 | pH_KCl | -0.73251 | 0.732511 | 10.51282 | 0.536572 |
| Dim.2 | Org_Carbon | 0.503215 | 0.503215 | 4.961335 | 0.253226 |
| Dim.2 | CEC | 0.698563 | 0.698563 | 9.560977 | 0.48799 |
| Dim.2 | Exch_Na | 0.54201 | 0.54201 | 5.755789 | 0.293774 |
| Dim.2 | Base_Saturation | -0.55917 | 0.559169 | 6.125999 | 0.31267 |
| Dim.2 | Exch_H | 0.698493 | 0.698493 | 9.559065 | 0.487893 |
| Dim.2 | Exch_Al | 0.78371 | 0.78371 | 12.03378 | 0.614202 |
| Dim.3 | Texture_Sand | -0.68084 | 0.680835 | 20.44347 | 0.463537 |
| Dim.3 | Texture_Silt | 0.630352 | 0.630352 | 17.52416 | 0.397344 |
| Dim.3 | Texture_Clay | 0.588988 | 0.588988 | 15.29972 | 0.346907 |
| Dim.3 | T_Nitrogen | -0.52626 | 0.526256 | 12.21419 | 0.276946 |
| Dim.4 | Carbon.Nitrogen | -0.61228 | 0.612282 | 26.01007 | 0.374889 |

**Table S9 | Final Weight Table for the final MDS Variables.**

Variables include Org_Carbon = organic carbon; CEC = cation exchange capacity; pH_H2O = soil pH measured in water (H₂O); P_Bray1 = Bray-1 extractable phosphorus; T_Nitrogen = total nitrogen

| Variable | PC | Loading | Abs Loading | Contribution | Cos2 | Wi |
| --- | --- | --- | --- | --- | --- | --- |
| CEC | Dim 2 | 0.699 | 0.699 | 9.56 | 0.488 | 0.219 |
| Org Carbon | Dim 2 | 0.503 | 0.503 | 0.503 | 0.253 | 0.114 |
| P Bray1 | Dim 1 | 0.579 | 0.579 | 0.579 | 0.336 | 0.105 |
| T Nitrogen | Dim 3 | -0.526 | 0.526 | 0.526 | 0.277 | 0.279 |
| pH H20 | Dim 2 | -0.795 | 0.795 | 0.795 | 0.633 | 0.284 |

**Table S10 | Visual Evaluation of Soil Structure (VESS) and soil structural quality (Sq) measurements.**

Measured across restoration treatments, planting types, species, and distances from the river edge. Restoration treatments comprised active enrichment planting beneath retained mature oil palms (MP + P) and active enrichment planting following mature palm removal (P Only), replicated across two river sites. Planting type refers to conspecific plantings, in which neighbouring seedlings were of the same species, and mixed plantings, in which neighbouring seedlings were of different species. Measurements were replicated across three enrichment-planted tree species (*Alstonia scholaris* (Pulai), *Peronema canescens* (Sungkai), and *Falcataria moluccana* (Sengon) and at different distances from the river edge. *Missing Sq depth* represents the depth of soil within the 0–25 cm VESS soil block that could not be assigned an Sq score. *VESS_0_25* represents the VESS score for the soil block, and *Sq health* represents the corresponding soil structural quality classification according to the VESS framework.

| Treatment | River Site | Plant ID | Planting | Species | Distance | Missing Sq depth | VESS_0_25 | Sq health |
| --- | --- | --- | --- | --- | --- | --- | --- | --- |
| MP + P | 1 | PC01 | Conspecific | Pulai | 32.7 | 0 | 3.14 | Poor / degraded |
| MP + P | 1 | PC02 | Conspecific | Pulai | 19.53 | 0 | 2.32 | Good / acceptable |
| MP + P | 1 | PC03 | Conspecific | Pulai | 9.34 | 0 | 2.94 | Good / acceptable |
| MP + P | 1 | PM01 | Mixed | Pulai | 6.14 | 0 | 2.4 | Good / acceptable |
| MP + P | 1 | PM02 | Mixed | Pulai | 11.73 | 0 | 4 | Poor / degraded |
| MP + P | 1 | PM03 | Mixed | Pulai | 8.26 | 15 | 2.4 | Good / acceptable |
| MP + P | 1 | SC01 | Conspecific | Sengon | 39.94 | 0 | 3.76 | Poor / degraded |
| MP + P | 1 | SC02 | Conspecific | Sengon | 13.62 | 0 | 3.5 | Poor / degraded |
| MP + P | 1 | SC03 | Conspecific | Sengon | 17.55 | 0 | 2.04 | Good / acceptable |
| MP + P | 1 | SM01 | Mixed | Sengon | 21.38 | 0 | 2.42 | Good / acceptable |
| MP + P | 1 | SM02 | Mixed | Sengon | 32.52 | 0 | 3.28 | Poor / degraded |
| MP + P | 1 | SM03 | Mixed | Sengon | 15.41 | 0 | 3.24 | Poor / degraded |
| MP + P | 1 | SUC01 | Conspecific | Sungkai | 21.62 | 0 | 3.1 | Poor / degraded |
| MP + P | 1 | SUC02 | Conspecific | Sungkai | 17 | 10 | 2.466667 | Good / acceptable |
| MP + P | 1 | SUC03 | Conspecific | Sungkai | 36.32 | 0 | 3.2 | Poor / degraded |
| MP + P | 1 | SUM01 | Mixed | Sungkai | 3.74 | 0 | 3.1 | Poor / degraded |
| MP + P | 1 | SUM02 | Mixed | Sungkai | 34.47 | 0 | 3.68 | Poor / degraded |
| MP + P | 1 | SUM03 | Mixed | Sungkai | 34.23 | 0 | 2.6 | Good / acceptable |
| MP + P | 2 | PCK01 | Conspecific | Pulai | 21.9 | 0 | 3.04 | Poor / degraded |
| MP + P | 2 | PCK02 | Conspecific | Pulai | 40.71 | 11 | 3.428571 | Poor / degraded |
| MP + P | 2 | PCK03 | Conspecific | Pulai | 43.31 | 0 | 3.68 | Poor / degraded |
| MP + P | 2 | PMK01 | Mixed | Pulai | 46.44 | 0 | 2.8 | Good / acceptable |
| MP + P | 2 | PMK02 | Mixed | Pulai | 33.68 | 0 | 3.12 | Poor / degraded |
| MP + P | 2 | PMK03 | Mixed | Pulai | 34.46 | 0 | 3.5 | Poor / degraded |
| MP + P | 2 | SCK01 | Conspecific | Sengon | 35.3 | 8 | 3.264706 | Poor / degraded |
| MP + P | 2 | SCK02 | Conspecific | Sengon | 34.31 | 7.5 | 2.942857 | Good / acceptable |
| MP + P | 2 | SCK03 | Conspecific | Sengon | 13.53 | 0 | 3.64 | Poor / degraded |
| MP + P | 2 | SMK01 | Mixed | Sengon | 11.83 | 0 | 2.98 | Good / acceptable |
| MP + P | 2 | SMK02 | Mixed | Sengon | 28.15 | 0 | 3.18 | Poor / degraded |
| MP + P | 2 | SMK03 | Mixed | Sengon | 16.55 | 0 | 3.02 | Poor / degraded |
| MP + P | 2 | SUCK01 | Conspecific | Sungkai | 33.96 | 0 | 2.64 | Good / acceptable |
| MP + P | 2 | SUCK02 | Conspecific | Sungkai | 17.5 | 0 | 3.44 | Poor / degraded |
| MP + P | 2 | SUCK03 | Conspecific | Sungkai | 39.01 | 0 | 2.6 | Good / acceptable |
| MP + P | 2 | SUMK01 | Mixed | Sungkai | 25.39 | 0 | 2.7 | Good / acceptable |
| MP + P | 2 | SUMK02 | Mixed | Sungkai | 21.66 | 0 | 3.72 | Poor / degraded |
| MP + P | 2 | SUMK03 | Mixed | Sungkai | 33.24 | 0 | 4.54 | Poor / degraded |
| P Only | 1 | PC05 | Conspecific | Pulai | 12.17 | 0 | 3.2 | Poor / degraded |
| P Only | 1 | PC06 | Conspecific | Pulai | 33.23 | 0 | 2.4 | Good / acceptable |
| P Only | 1 | PC07 | Conspecific | Pulai | 15.15 | 0 | 3.04 | Poor / degraded |
| P Only | 1 | PM07 | Mixed | Pulai | 19.65 | 0 | 3.84 | Poor / degraded |
| P Only | 1 | PM08 | Mixed | Pulai | 25.97 | 0 | 3.16 | Poor / degraded |
| P Only | 1 | PM09 | Mixed | Pulai | 25.25 | 0 | 3.2 | Poor / degraded |
| P Only | 1 | SC05 | Conspecific | Sengon | 9.31 | 0 | 3.2 | Poor / degraded |
| P Only | 1 | SC06 | Conspecific | Sengon | 8.95 | 2 | 2.652174 | Good / acceptable |
| P Only | 1 | SC07 | Conspecific | Sengon | 16.42 | 0 | 2.94 | Good / acceptable |
| P Only | 1 | SM04 | Mixed | Sengon | 23.75 | 0 | 2.88 | Good / acceptable |
| P Only | 1 | SM07 | Mixed | Sengon | 31.38 | 0 | 2.6 | Good / acceptable |
| P Only | 1 | SM08 | Mixed | Sengon | 17.46 | 0 | 1.88 | Good / acceptable |
| P Only | 1 | SUC07 | Conspecific | Sungkai | 29.26 | 0 | 2.28 | Good / acceptable |
| P Only | 1 | SUC08 | Conspecific | Sungkai | 36.45 | 0 | 3.48 | Poor / degraded |
| P Only | 1 | SUC09 | Conspecific | Sungkai | 13.31 | 0 | 2.24 | Good / acceptable |
| P Only | 1 | SUM05 | Mixed | Sungkai | 20.7 | 0 | 2.84 | Good / acceptable |
| P Only | 1 | SUM07 | Mixed | Sungkai | 10.51 | 0 | 3.04 | Poor / degraded |
| P Only | 1 | SUM10 | Mixed | Sungkai | 8.14 | 0 | 3.24 | Poor / degraded |
| P Only | 2 | PCK04 | Conspecific | Pulai | 21.68 | 0 | 3.56 | Poor / degraded |
| P Only | 2 | PCK05 | Conspecific | Pulai | 24.81 | 0 | 3.66 | Poor / degraded |
| P Only | 2 | PCK06 | Conspecific | Pulai | 30.46 | 0 | 2.66 | Good / acceptable |
| P Only | 2 | PMK05 | Mixed | Pulai | 36.37 | 0 | 3.32 | Poor / degraded |
| P Only | 2 | PMK06 | Mixed | Pulai | 26.12 | 0 | 3.28 | Poor / degraded |
| P Only | 2 | PMK07 | Mixed | Pulai | 21.86 | 0 | 3 | Poor / degraded |
| P Only | 2 | SCK04 | Conspecific | Sengon | 40.1 | 0 | 3.24 | Poor / degraded |
| P Only | 2 | SCK05 | Conspecific | Sengon | 17.9 | 0 | 2.4 | Good / acceptable |
| P Only | 2 | SCK06 | Conspecific | Sengon | 12.67 | 0 | 3.62 | Poor / degraded |
| P Only | 2 | SMK04 | Mixed | Sengon | 38.04 | 0 | 2.4 | Good / acceptable |
| P Only | 2 | SMK05 | Mixed | Sengon | 29.13 | 0 | 3.26 | Poor / degraded |
| P Only | 2 | SMK06 | Mixed | Sengon | 31.74 | 0 | 2.92 | Good / acceptable |
| P Only | 2 | SUCK04 | Conspecific | Sungkai | 35.57 | 0 | 4.08 | Poor / degraded |
| P Only | 2 | SUCK05 | Conspecific | Sungkai | 13.77 | 0 | 3.74 | Poor / degraded |
| P Only | 2 | SUCK06 | Conspecific | Sungkai | 37.89 | 0 | 2.32 | Good / acceptable |
| P Only | 2 | SUMK04 | Mixed | Sungkai | 38.57 | 0 | 3.88 | Poor / degraded |
| P Only | 2 | SUMK05 | Mixed | Sungkai | 35.15 | 0 | 2.64 | Good / acceptable |
| P Only | 2 | SUMK06 | Mixed | Sungkai | 35.69 | 0 | 3.16 | Poor / degraded |
